# Machine Learning Reveals Distinct Immunogenic Signatures of Th1 Imprinting in ART-Treated Individuals with HIV Following Repeated SARS-CoV-2 Vaccination

**DOI:** 10.1101/2025.03.18.643769

**Authors:** Chapin S. Korosec, Jessica M. Conway, Vitaliy A. Matveev, Mario Ostrowski, Jane M. Heffernan, Mohammad Sajjad Ghaemi

**Affiliations:** Modelling Infection and Immunity Lab, Mathematics and Statistics, York University, 4700 Keele St, Toronto, M3J 1P3, ON, Canada; Centre for Disease Modelling, Mathematics and Statistics, York University, 4700 Keele St, Toronto, M3J 1P3, ON, Canada; Department of Mathematics, Pennsylvania State University, 201 Old Main, University Park, University Park, 16802, Pennsylvania, United States of America; Department of Medicine, University of Toronto, 27 King’s College Cir, Toronto, M5S 1A8, Ontario, Canada; Department of Immunology, University of Toronto, 1 King’s College Cir, Toronto, M5S 1A8, Ontario, Canada; Keenan Research Centre for Biomedical Science, St. Michael’s Hospital, Unity Health, Toronto, M5B 1W8, Ontario, Canada; Digital Technologies Research Centre, National Research Council Canada, 222 College Street, Toronto, M5T 3J1, ON, Canada

**Keywords:** SARS-CoV-2 vaccination, HIV, Antiretroviral therapy (ART), Machine Learning (ML), Random forests (RFs), Synthetic data, Immunology, Vaccinology, Adaptive immunity, Th1 imprinting, Personalized vaccination strategies, Biomarker classification, Immune dysregulation

## Abstract

The human immune system is intrinsically variable and remarkably diverse across a population. The immune response to antigens is driven by a complex interplay of time-dependent interdependencies across components of the immune system. After repeated vaccination, the humoral and cellular arms of the immune response display highly heterogeneous dynamics, further complicating the attribution of a phenotypic outcome to specific immune system components. We employ a random forest (RF) approach to classify informative differences in immunogenicity between older people living with HIV (PLWH) on ART and an age-matched control group who received up to five SARS-CoV-2 vaccinations over **104** weeks. RFs identify immunological variables of importance, interpreted as evidence for Th1 imprinting, and suggest novel distinguishing immune features, such as saliva-based antibody screening, as promising diagnostic features towards classifying responses (whereas serum IgG is not). Additionally, we implement supervised and unsupervised *Machine Learning* methods to produce physiologically accurate synthetic datasets that conform to the statistical distribution of the original immunological data, thus enabling further data-driven hypothesis testing and model validation. Our results highlight the effectiveness of RFs in utilizing informative immune feature interdependencies for classification tasks and suggests broad impacts of ML applications for personalized vaccination strategies among high-risk populations.

## 1 Introduction

HIV remains a global health burden with approximately 40 million current infections (People living with HIV, PLWH) and over 40 million deaths to date [1]. Given the current global trends in new infections, millions of new infections are expected by 2050 [2]. Antiretroviral therapy (ART) has significantly improved life expectancy and overall health outcomes for PLWH [3, 4]. However, effective HIV viral load suppression may not lead to full recovery [5–8].

Vaccines are the premier prophylactic public health intervention to reduce infectious disease severity and morbidity. Longitudinal vaccine-induced immunogenicity against vaccine-preventable infections has been studied in the context of immunodeficiencies in ART-suppressed PLWH for different pathogens and vaccine types [9, 10]. Vaccine-elicited responses in ART-suppressed PLWH have often been found to be inferior to HIV-negative control groups, although the gap is often diminished following a multidose vaccination regimen [11]. Studies of variations in the humoral and cellular immune responses from repeated vaccination in PLWH and non-infected age-matched controls may shed light on the effects of persistent immune activation due to ART-suppressed HIV infection on the adaptive immune response dynamics. This knowledge could inform customized vaccination strategies [12], aid in the development of adjuvant therapies [13], and reduce the risk of severe outcomes [14] in the PLWH immunologically vulnerable group.

Induction of the adaptive immune response to vaccines and pathogens requires the activation and proliferation of CD4+ T-helper cells that subsequently activate and generate cellular and humoral immunity [15, 16]. The cellular and humoral components of the human immune system can be highly variable across a population, and they can exhibit complex time-sensitive interdependent immune responses to antigenic perturbations [17]. Among PLWH on ART, ART adherence and immune status (CD4 immune responder (IR) versus immune non-responder (INR)) may have an influence on the durability of humoral and cellular responses to SARS-CoV-2 vaccinations [18]. For example, humoral outcomes among PLWH with undetectable plasma HIV viral loads and high CD4 counts have been found to be similar to HIV-negative controls for the ChAdOx1 nCoV-19 vaccine [19], heterogeneous multidose vaccine regimens [7, 20], and mRNA-based vaccines [21, 22], however, PLWH classified as INR display CD8 and CD4 responses that significantly differ from controls [7, 18]. Given enough data, machine learning (ML) algorithms can capture and learn the immune signatures of PLWH versus a control group, and furthermore, may identify which variables are particularly informative or non-informative in distinguishing each individual’s immune class. An advantage of ML approaches is that they can reveal complex nonlinear relationships between cellular and humoral immune responses without requiring explicit assumptions about the nature of the immunological feature relationships [23]. The motivation for employing machine learning approaches is to create an advanced tool capable of processing complex datasets at the individual level. We aim to leverage machine learning to classify individuals with a defined probability while offering clear, data-driven explanations for each classification decision based on their personal data. With a successfully trained ML tool, consistently *misclassified* PLWH may represent individuals with atypical immune responses to HIV. These responses could include unusually effective viral suppression, the presence of unique genetic factors such as in elite controllers, or intriguing immunological variations in treatment responses. Conversely, age-matched HIV-negative individuals consistently *misclassified* as HIV-positive exhibit vaccine-elicited immune profiles resembling those of HIV-positive individuals. Such profiles could arise due to other previously unknown comorbidities, including chronic inflammation, autoimmune diseases, or concurrent infections. Machine learning offers a powerful framework for personalized vaccination strategies in immunology by enabling the classification of individuals based on complex immune signatures while providing interpretable insights into atypical responses. By identifying misclassified cases, we can uncover novel immunological variations, including unique host factors in PLWH and previously unrecognized comorbidities in HIV-negative individuals, ultimately refining precision medicine approaches for vaccine response assessment.

In the following, we investigate heterogeneity in the immune response from SARS-CoV-2 vaccine-elicited immunological outcomes among ART-suppressed PLWH who received up to five doses of the SARS-CoV-2 vaccines over 104 weeks. The HIV-specific responses are classified against an age-matched control group, thereby controlling for known age-related immunological variations and dysregulations [24]. The dataset that we study is an extended dataset from Matveev *et al.* [7]. We find that Random Forest, which is capable of learning complex nonlinear interdependencies between immunological features that traditional statistical and mechanistic models may have missed or are incapable of learning [25], is able to distinguish between the PLWH and the age-matched control immune responses with near-perfect accuracy, given particular biomarkers. Our findings reveal that there are informative and non-informative immunogenic features implicated in the adaptive immune response of PLWH; informative features are determined by the random forest to be very important in classifying immune responses, while non-informative immune features provide little information towards classifying the immune responses. For instance, while saliva and serum IgG were non-informative, saliva IgA emerged as a critical indicator in identifying HIV-modulated immunogenic responses; further, cytokine and saliva IgA features in combination are found to form an optimally stratified feature subset leading to equivalent performance metrics as the full dataset.

Finally, we extend our study to generate synthethic datasets that reproduce the local and global characteristics of the human-based dataset. We employ both supervised and unsupervised synthetic data generation methods such as multivariate normal, Gaussian mixture models, synthetic minority oversampling technique, and K-nearest neighbors. We demonstrate that the preservation of local or global data characteristics depends on the synthetic data generation method. Finally, we assess how well each synthetic dataset performs with RF classification and discuss future considerations.

## Results

To characterize the immunogenic responses elicited by repeated SARS-CoV-2 vaccinations in ART-treated individuals with HIV (PLWH) versus an age-matched control group, we employed a combination of unsupervised and supervised ML methods. Principal Component Analysis (PCA) and Linear Discriminant Analysis (LDA) were utilized to explore data clustering and class separability, respectively. A Random Forest (RF) classifier was then implemented to identify complex, nonlinear immunogenic signatures that differentiate the groups, while feature importance analyses indicated key immunological contributors to these differences. t-SNE-based correlation network visualization was then employed to enhance our understanding of the feature interrelationships. These approaches collectively provided a comprehensive framework for uncovering meaningful immunological patterns and for classifying responses with high accuracy. Finally, we utilized various supervised and unsupervised ML methods to accurately capture the local and global data characteristics, and demonstrate that synthetic data can reproduce the clustering and RF classification behaviour as the original dataset.

### Study data timeline, participants

Our data is an extended dataset from our previous study [7] to include doses 4 and 5 of COVID-19 vaccination. In total, 91 participants were recruited into the study – 23 HIV-negative individuals and 68 PLWH on ART. The timeline of the clinical study is shown in Figure 1A. Study participants were given 5 vaccine doses, and biomarker measurements are made in each study interval, see Methods and [7] for details. The full dataset used in this work is comprised of 63 immune features, consisting of serum and saliva IgG, saliva IgA, IFNg and IL2-producing T cells, and dual-responding IL2/IFNg cells, CD4/CD8 ratio, virus neutralization, and ACE2 displacement, drawn from individuals throughout the course of their multivaccine regimen across the study timeline (Figure 1A). Figure 1B plots the raw IgG RBD data.

**Fig. 1.**
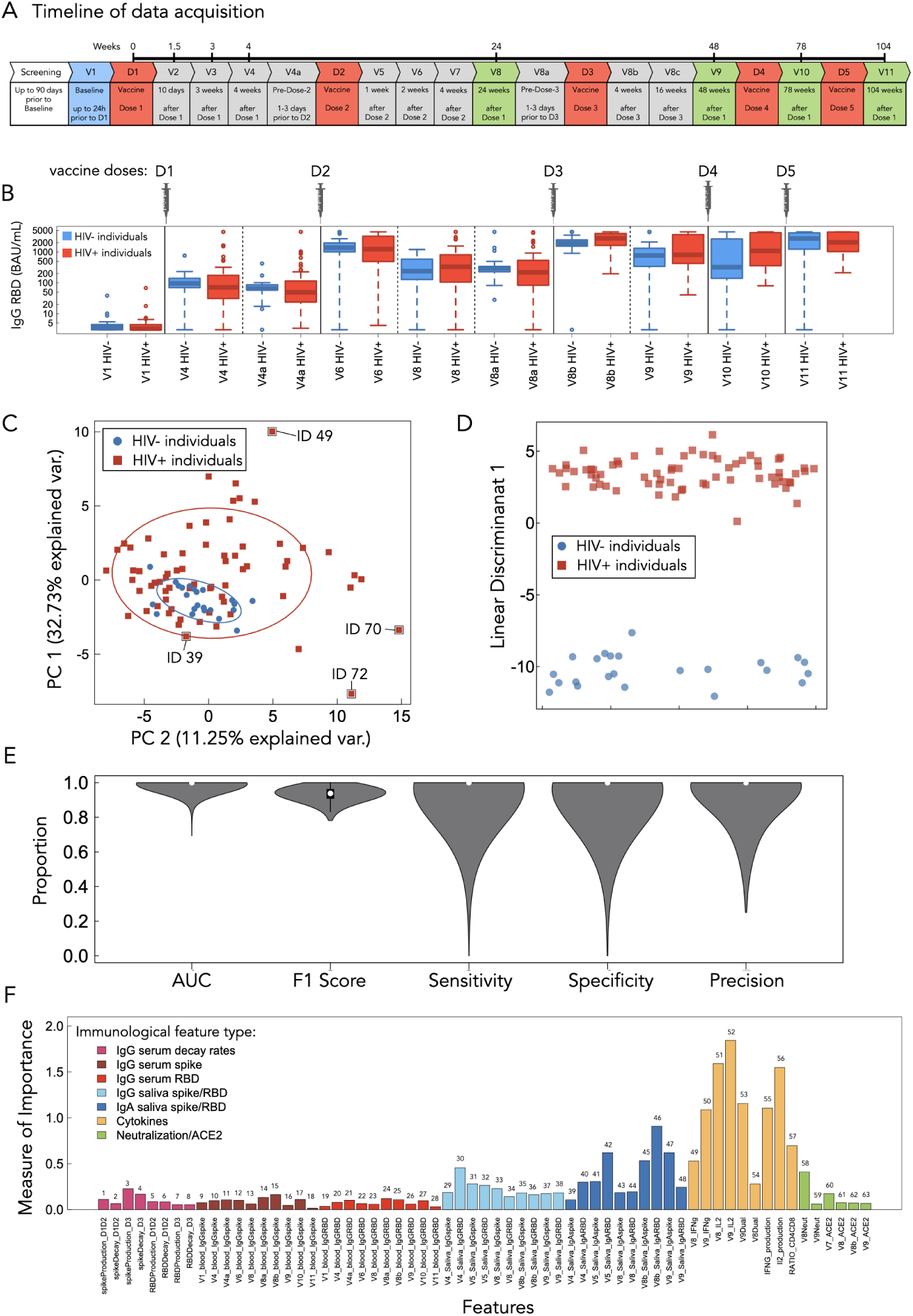
Dataset overview and RF performance metrics. A) Timeline of study and data acquisition. B) IgG RBD features for the HIV- and HIV+ classes for doses 1 through 5. C) PCA with the first two largest components shows that significant source of variation in the data is not driven by the HIV+ group. D) The first largest component (LD1) of LDA reveals the existence of HIV-specific signatures that are not distinguishable through the unsupervised PCA. E) The performance metrics—AUC-ROC, F1 Score, sensitivity, specificity, and precision—demonstrate the strong predictive power of the RF model. F) The Measure of importance, as assessed by the the RF algorithm is shown for all 63 features. Features are grouped by immunological type and colour coded.

### SARS-CoV-2 vaccine responses of PLWH and control can be classified with near-perfect performance

PCA is an unsupervised method that can be used to reduce the dimensionality of high-dimensional data by identifying the directions (principal components) that capture the most variance in the dataset. PCA can thus be used to learn whether vaccine-ellicited immune responses among PLWH versus the age-matched control differentiate the classes effectively. The PCA results do not show clear separation between the groups (Figure 1C). However, we observe that HIV-individuals subcluster within the HIV+ cluster (Figure 1C). Additionally, we find that PLWH exhibit a broader spread along both principal components 1 and 2 (PC1 and PC2, respectively), evidenced by the 95% confidence ellipses. We include labels for outlier study participant IDs 49, 70, and 72; we interpret outliers as individuals who may have distinctive immune responses.

In order to maximize class separation and check for separability, Linear Discriminant Analysis (LDA), a supervised and linear dimensionality reduction technique that focuses on maximizing the separation between predefined classes, was employed. The LDA results using the first linear discriminant (LD1) are shown in Figure 1D and present a clear separation between the HIV- and HIV+ groups. We thus conclude that the immunological features included in our study can be used to discriminate between the HIV- and HIV+ classes.

While PCA and LDA provide an intuitive visualization of data to explore class separability, these methods are not able to capture the complex inter-dependencies between features. However, the class separability pointed to by LDA suggests that a more advanced algorithm may be able to learn unique immunological signatures that capture the PLWH immune response that distinguishes it from the control population. We therefore used a random forest (RF) classifier to characterize the complex nonlinear signatures differentiating HIV+ and HIV-immunological responses in order to enhance predictive accuracy. This can also provide mechanistic insights on the underlying immunological trends. The details of the RF implementation can be found in Methods.

The RF models achieved a perfect median AUC-ROC (area under the receiver operator curve; measures a model’s ability to distinguish between classes, evaluating performance across all classification thresholds) of 1.0 (Figure 1E). AUC-ROC reflects the model’s ability to distinguish between the HIV positive and negative classes across all possible classification thresholds. The AUC-ROC median value of 1.0 suggests that the RF classifier has excellent overall classification performance on our immunological dataset.

The RF model’s median F1 score (the harmonic mean of precision and recall, balancing false positives and false negatives) was found to be 0.94 (Figure 1E). This reveals a trade-off between precision (the proportion of correctly identified positive instances) and recall (the proportion of actual positives correctly identified) and suggests that there may be a moderate number of false positives (affecting precision) or false negatives (affecting recall) in the predictions. We report median values of 1.0 for sensitivity, precision, and specificity (Figure 1E), which all suggest that the model often achieves near-perfect performance, but considerable variance around these median values implies that this performance is not consistent across all data folds.

AUC-ROC and F1 Score performance measures from RF trained on individuals with randomized labels are all found to be ∼ 0.5 on median (Figure S1). To further explore RF classification heterogeneity for each individual, as well as the likelihood of HIV+ classification across all models, we provide all probability distributions among all IDs in Figure S2A,B.

We conclude that our implementation of RF is generally effective at classifying vaccine-elicited immunogenic responses among PLWH vs the age-matched control group. Further exploration of the RF results and the significance of the outliers is discussed below.

### RF reveals T-cell responses to be most important features in distinguishing the vaccine-elicited immune response between PLWH and the age-matched HIV-control group

Successfully mounting, and maintaining, vaccine-elicited immunity involves a myriad of immunological components (for a good review see ref. [15]). Immunodeficient states, such as those that can be found in ART-suppressed PLWH, can have complex dysregulated immune signatures – where particular components of the immune response can be dysregulated more than others. In our feature space, we are therefore interested in using the RF algorithm to suggest which immune components are informative in distinguishing the PLWH SARS-CoV-2 vaccine-ellicited responses from the age-matched controls. Such insights may be useful towards forming mechanistic hypotheses for how ART-suppressed HIV infection affects adaptive immune system dynamics.

To assess how each feature contributes to RF model accuracy we computed the feature importance (see Method for details). We plot the importance estimates in Figure 1F for all 63 immune features. Features with higher importance scores have a more significant impact on improving model predictions in decision tree splits that underlie the random forest, suggesting they capture significant information that can differentiate the classes. Our analysis indicates that the frequencies of SARS-CoV-2 spike-specific T cells measured by cytokine production are generally the most important features, with the three interleukin-2 (IL2) features (Visit 8,9, and IL2 production rate) having the highest importance over all other features. Conversely, the serum-based humoral features all rank as least important, suggesting information contained within these immune features is redundant towards classification. Below, the importance rankings (Figure 1F) will be used in forward and reverse ablation analyses to determine if there exists a minimally stratified combination of features that leads to optimal RF performance.

### tSNE reveals that important features tend to form clusters

We use t-stochastic neighbour embedding (t-SNE) to visualize the structure of our feature dataset. Figure 2A presents the correlation network of the dataset using all 63 biomarkers where circle size represents the corresponding biomarker RF importance. Here, we learn that IgA, cytokines, and IgG biomarkers tend to cluster together, but the cluster densities may increase if there are more important biomarkers within that class. We also learn that biomarkers that are determined to be important (e.g., IL2 and IFNs) tend to form distinct groupings, suggesting correlated expression patterns, while features that are determined unimportant (e.g. spike and RBD IgG) disperse, suggesting weaker correlations among these features. Figure 2 panels B and C display the same t-SNE layout as panel A, however, only significant edges connecting to a cytokine feature are displayed, where colour is now used to illustrate the clear and opposing RF feature weight trends found for HIV- (panel B) and HIV+ (panel C) individuals, respectively. From this we can see that there are clearly opposing RF-weights assigned to the cytokines, depending on whether the individuals are PLWH or the age-matched control. The t-SNE layout reveals that the most important features identified by the RF algorithm form distinct clusters, suggesting that these features not only drive the predictive power of the RF model but also represent well-defined biologically meaningful patterns in the data (Figure 2 A), with the RF oppositely weighting key features to distinguish the classes (Figure 2B,C). We next plot the full distributions for all feature weights across all folds to visualize the distributions of RF weights. The feature weight distributions from all folds from all (*n* = 23) control individuals and from all (*n* = 68) PLWH are shown in Figure 2D, and Figure 2E, respectively. We see more clearly that classification of the control class is highly dependent on the cytokine measures, which have negative weights on median. There is very little weight placed on any of the other features except V8B IgA RBD saliva, whose weight distributions deviate below 0 on median. Conversely, for PLWH we find that cytokine features are positively weighted (Figure 2E, yellow features). We also find the V8B IgA RBD saliva values to be positively weighted, opposite that of the control. While the median weights for all other features are ∼0, we observe significantly more dispersion in median values for the PLWH compared to the control (Figure 2E, all non-yellow features).

**Fig. 2.**
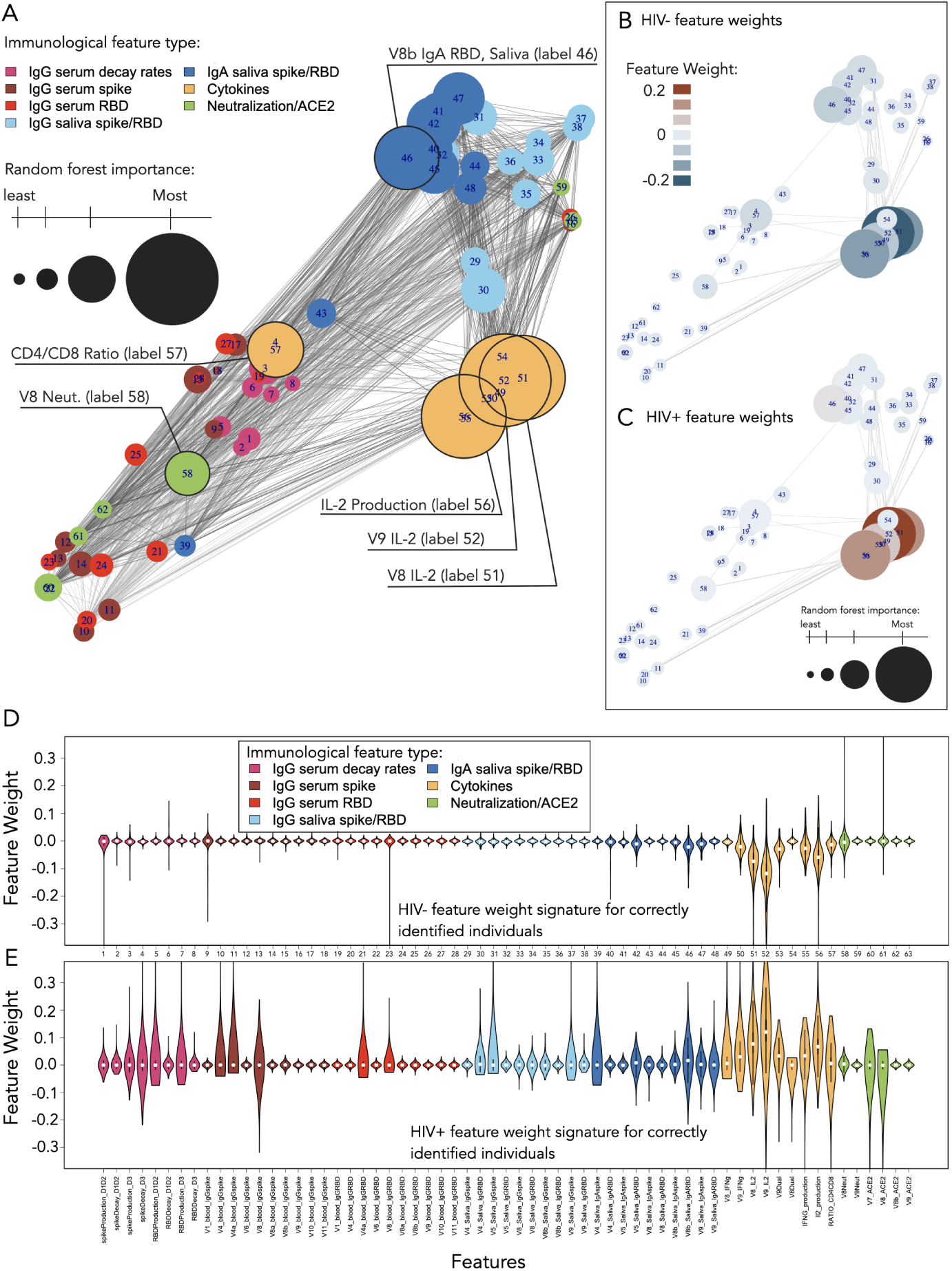
Correlation network and feature weight distributions. A) A correlation network of the 63 immunological features is visualized using t-SNE applied to the adjacency matrix to calculate the layout. Features are coloured by type, and node sizes scale relative to the importance measures from Figure 1F. For visualization, each feature is represented as a node. An edge is shown between a pair of nodes when their correlation is statistically significant (*p-values* < 0.05). B,C) The subset of network connections with statistically significant correlations, that further depend on the cytokine features, is shown. Here, the colour scale corresponds to the mean feature weights for HIV- (panel B) and HIV+ (panel C) individuals. D,E) the distributions of RF feature weights computed across all models is shown for the individuals who are correctly classified more than 95% of the time. Median cytokine feature weights (yellow) clearly form opposite trends when comparing HIV- (panel D) and HIV+ (panel E) RF results.

Figure 2D shows that the control class is relatively homogeneous (most feature weights are close to 0) and that only cytokines are needed to distinguish HIV-individuals from the HIV+ class. Figure 2E shows significant RF feature weight variance for the HIV+ class for serum and saliva IgG biomarkers in addition to heavily weighted cytokine features, suggesting that many more features are needed, potentially in combination, for correct classification. The significant dispersion found in 2E suggests heterogeneity in disease pathology and immune responses across PLWH, indicating that, due to the high variability and complexity of the HIV+ immune response, a diverse set of trees in the forest are required to accurately classify instances of the HIV+ class, where each tree may rely on different combinations of features. Therefore, we next study the RF performance under various ablation procedures to determine a minimum stratified set of features for optimal classification. We also explore sensitivity to RF performance with various combinations of features by their clinical type.

### Ablation analysis reveals optimally-stratified feature subset for classification

Ablation analysis identifies which features are essential for classification and which are redundant or irrelevant. Identifying a minimally stratified dataset reveals the key immunological markers (e.g., specific cytokines or antibodies) that distinguish HIV+ from HIV-individuals and thus, can enhance understanding of the biological mechanisms driving the immune response in HIV infection. Further, the immunological dataset may include noisy features. Identifying and removing these features can improve model clarity and reduce overfitting.

To identify the minimal set, we perform both forward and reverse ablation (see Methods). Results are displayed in Figure 3A. Forward ablation (blue hollow squares) reveals that just the top two features (V8 and V9 IL2 measures) can be used to produce a median AUC-ROC just below 1.0. Conversely, reverse ablation results in a monotonic decrease in median AUC-ROC (yellow hollow triangles) down to a minimum median of ∼0.65. The final two features identified via reverse ablation are the visit 11 blood IgG spike and visit 11 blood IgG RBD features.

**Fig. 3.**
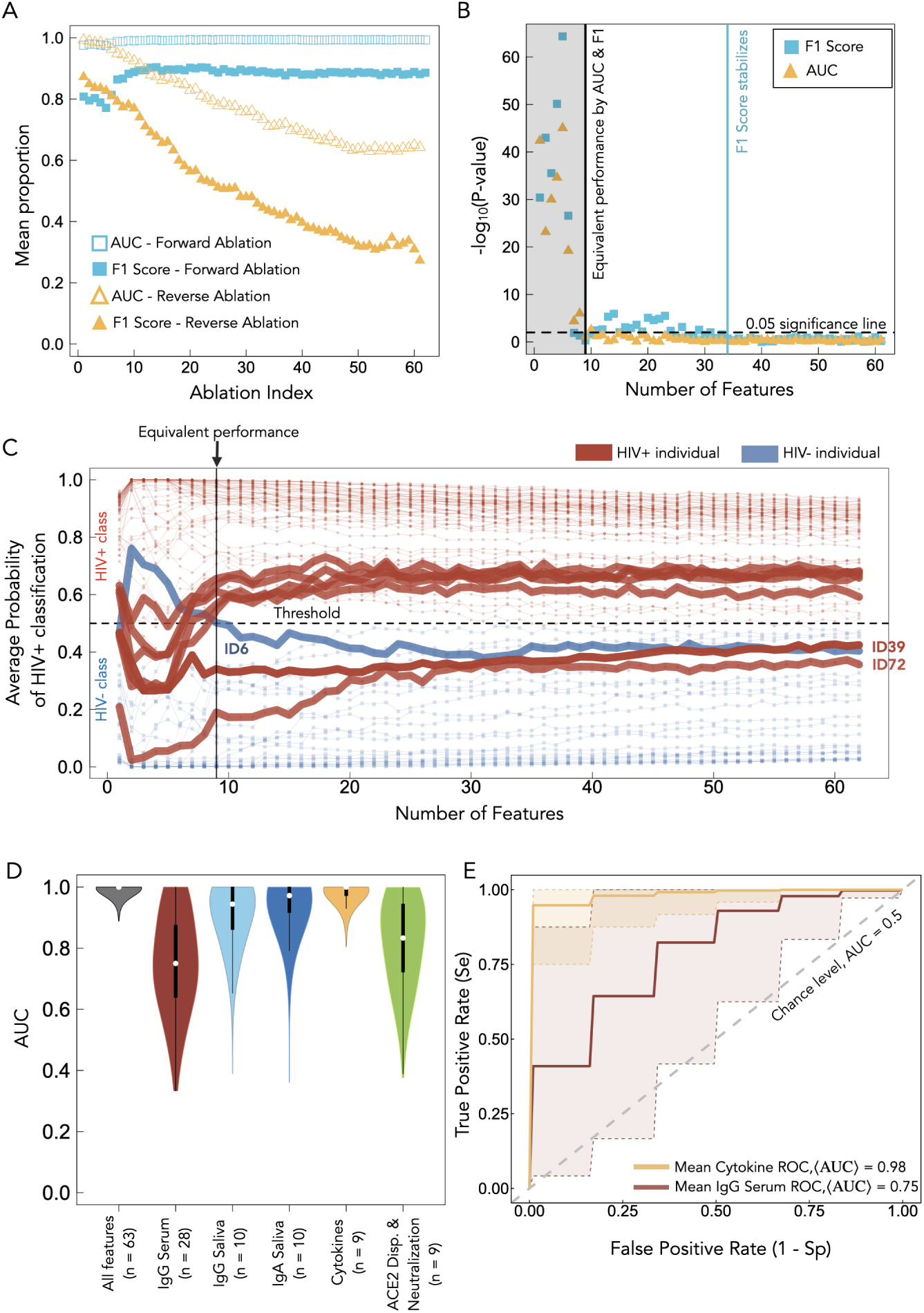
Ablation analyses and model reduction. A) Mean AUC-ROC and F1 score are shown as a function of the ablation index for the forward and reverse ablation analyses. B) P-values from Wilcoxon signed-rank test performed between the full-feature RF model and each of the performance-measure distributions from the forward ablation procedure are shown. C) Average probability trajectories for all 91 individuals as a function of number of features from the forward ablation algorithm. Multiple individuals whose probability of classification crosses over the threshold at index 9 are shown at the equivalent performance mark (solid line). D) AUC-ROC distributions of RF performance are shown for RF models trained on features in isolation by their clinical type. Here, *n* refers to the number of features of each type. E) Mean ROC curves generated using predictions from the test set for RF models trained on cytokine (yellow), and IgG serum (red) data types are shown. Shaded regions correspond to 95% confidence intervals are highlighted by a dashed line.

Note that the forward ablation F1-score follows a non-monotonic relationship as a function of forward ablation index (blue filled squares). The top two features produce an F1 Score of ∼ 0.8 (Forward ablation index 1, first blue square in Figure 3A), but the F1 Score decreases to a minimum of ∼ 0.76 by forward ablation index 6 before increasing up to a plateau of ∼0.9 as additional features are added. As a function of reverse-ablation index, the F1 Scores (yellow-filled triangles) decrease, as expected, to a minimum of ∼0.3. Thus, the least two important features lead to F1 Scores of 0.3 and and AUC-ROC of 0.7 (Figure 3A solid and hollow yellow triangles, respectively).

Fig 3B demonstrates the quality of the identified minimal set – a Wilcoxon test is used to compare the AUC-ROC and F1 Score distributions from the forward ablation algorithm to the full 63 feature distributions (shown in Fig 1E). Considering a p-value of 0.05 to be the significance threshold, we find the top 9 feature performance to be statistically similar to the complete 63 feature dataset, with both the AUC-ROC and F1 scores used for comparison. Thus, these 9 features may constitute an optimally-stratified feature set for classifying SARS-CoV-2 vaccine immune responses between HIV+ and HIV-immune statuses. Note that following the ninth forward ablation index, the AUC-ROC remains under the 0.05 significance line (Figure 3B, yellow triangles), while the F1 Score-associated p-value increases to above significance and does not settle to below significance until the 34th feature is added (Figure 3B, blue squares). Therefore, the addition of more features beyond the optimally stratified set of features adds redundant information and can lead to a tendency to misclassify some individuals. In decreasing order of importance, the 9 features in the optimal set are: Visit 9 IL2, Visit 8 IL2, the IL2 production rate, Visit 9 Dual responding cells, the IFNg production rate, Visit 9 IFNg, Visit 8b Saliva IgA RBD, the CD4/CD8 ratio, and visit 5 saliva IgA RBD. For clarity, the IL2 and IFNg production rates refer to the rate of change of the increasing frequencies of cytokine-producing T cells, as a measure of spike-specific T-cells, during the course of the booster series, calculated in our previous work [7]. Thus, seven of the top nine features are cytokine-based while two are saliva IgA RBD-based features.

### Saliva humoral responses are good predictors of PLWH SARS-CoV-2 vaccine immunogenecity

Figure 3D shows the AUC-ROC distributions of RF performance for RF models trained according to the characteristics of their clinical type, ie, serum IgG, saliva IgG, saliva IgA, cytokines and neutralization / ACE2 displacement. Figure 3E displays the mean ROC curves with 95% confidence intervals for cytokines (best performing feature family) and serum IgG (worst performing feature family). In Fig S5 we provide the accompanying mean PR plot for the cytokine and serum data. These results show that the RF model trained on cytokine features provides a near-perfect classifer. The IgA and IgG saliva features then take second and third place, in terms of AUC-ROC performance, with median AUC-ROC values of 0.98 and 0.95, respectively. Classification performance then drops when using ACE2 displacement/Neutralization and IgG serum features to median values of 0.83 and 0.75, respectively. The IgG serum features thus provide a near-baseline classifier. There is large dispersion (standard deviation) in AUC’s found for the last two clinical feature classes of 0.14 and 0.16, respectively. The significant biological ramifications of this result, specifically that mucosal IgG and IgA appear to be highly informative while serum IgG is not, are highlighted in the discussion in the context of known HIV-related mucosal immune dysregulatory mechanisms.

### Misclassified individuals suggest atypical vaccine immune responses

The identification of individuals whose immunological responses lead to classification errors may suggest closer follow up by clinicians. For example, consistently misclassified PLWH may represent atypical immune responses to HIV, such as unusually effective viral suppression, unique genetic factors (e.g., elite controllers), or interesting immunological variations in treatment responses. Conversely, age-matched HIV-individuals consistently misclassified as HIV+ might have immune profiles resembling those of HIV-positive individuals due to other comorbidities, such as chronic inflammation, autoimmune diseases, or infections. To further investigate the non-monotonic F1 Scores, we identify which individuals are contributing to the initial decline in F1 Score shown in Figure 3A. Fig 3C provides the average probability of HIV+ classification for all individuals as a function of forward ablation index. We highlight a number of trajectories that by index 9 (the equivalent performance index) cross the classification threshold towards the correct region, and note that after index 9 all individual predictions, on average, remain in their respective classification region. The visualization provided in Fig 3C provides a deeper intuition of the model performance metrics displayed in Fig 3A,B. IDs 39 and 72, two PLWH, are the only IDs to remain in the incorrect region beyond forward ablation index 9, and remain so when all features are present. IDs 39 and 72 are misclassified in 95% and 85% of the iterations, respectively, while there are seven individuals misclassified 10-50% of the time. The remaining 82 individuals are classified correctly in nearly 100% of the tests. Figure S2 provides an additional visualization displaying the probabilities of HIV classification for all individuals in the study, as well as ID 39’s unique immunological weight signature which appears similar to the HIV-control signature.

Figure S2B displays the HIV+ classification probabilities for all 91 individuals. In Figure S2C we include the individual feature weight distributions, across all RF models, for ID39 who is an HIV+ individual misclassified 95% of the time. ID39’s cytokine feature weight signature matches the HIV-feature signature shown in Figure 2D. Further, an example of a single training landscape from a randomly selected RF model is shown in Figure S3, with two arbitrary study participant ID examples illustrated in Figure S4. Altogether, these results suggest there may be redundant features present that may mislead the RF algorithm, or there are outlying individuals with unique signatures (e.g. ID39) that result in the RF algorithm placing significant weight on non-cytokine features.

### Synthetic data provides physiologically-accurate representation of immunological data

We implemented Gaussian Mixture Models (GMM), Multivariate Normal distributions (MVN), synthetic oversampling technique (SMOTE), and K-nearest neighbours (KNN) methods in order to generate synthetic data. We ensure that the generated data matches the original dataset in size for all iterations: 64 features total, comprising 23 HIV-individuals, and 68 PLWH. Detailed descriptions of all ML algorithms performed can be found in Methods.

GMM leads to the lowest mean Kullback leibler divergence (KLD) and second-lowest KLD variance, followed by the MVN approach (Figure S6). By KLD metrics, the worst performance is found to be SMOTE, closely followed by KNN (for all three k values examined). Figure 4A shows each calculated KLD value for all 63 features for the GMM approach, where a broad distribution of KLD values as a function of feature type is revealed. Figure 4A prompted further analysis to determine whether some synthetic generating techniques are better for specific features than others. In Figure S7 we provide a t-SNE plot, with the same layout as in Figure 2A, where colour corresponds to the synthetic data generating technique that produced the lowest KLD for each feature. We find no discernable relationship between synthetic generation method and feature cluster behaviour.

**Fig. 4.**
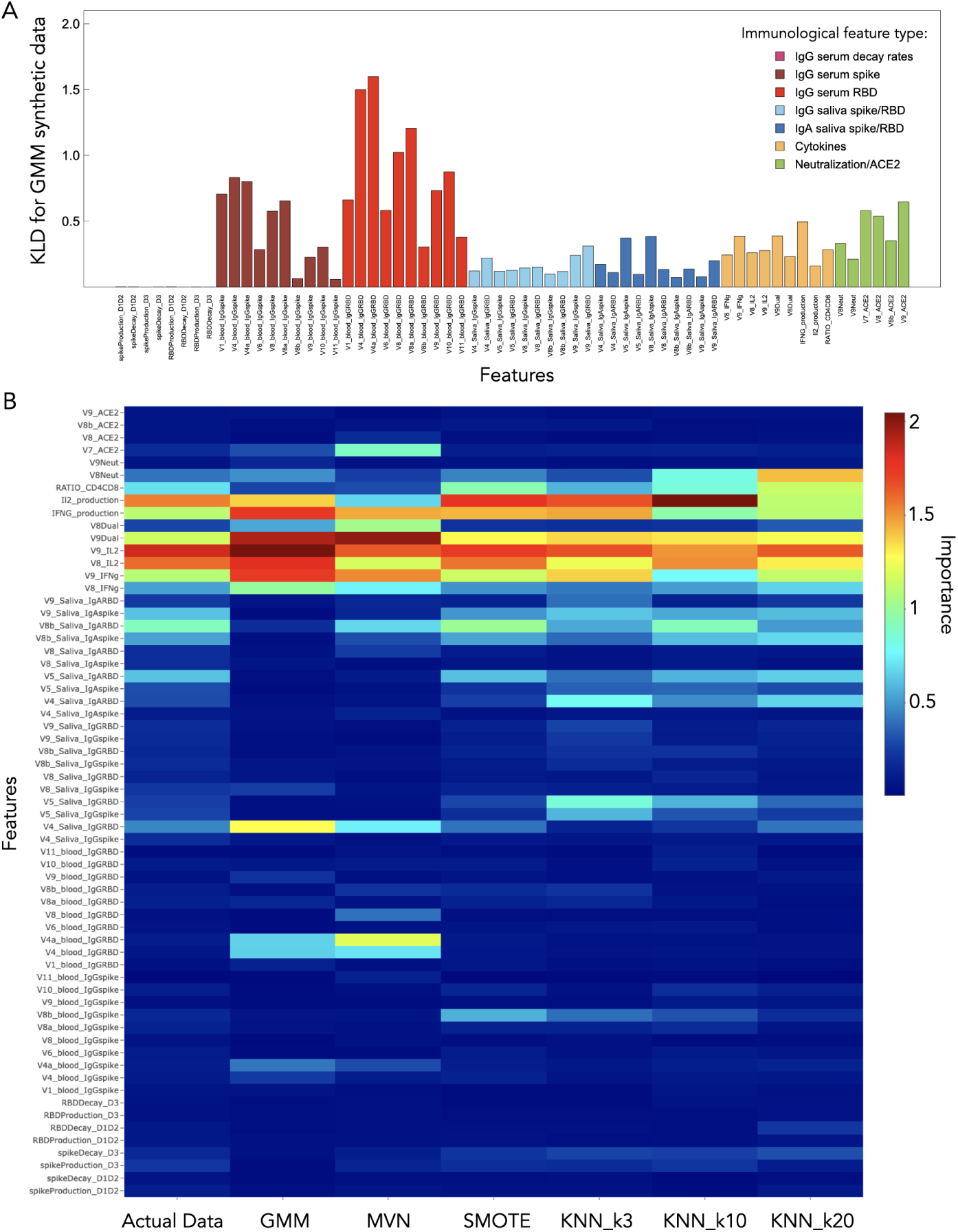
Synthetic data analysis. A) KLD is shown for each feature for the GMM synthetic data, which provided the lowest mean KLD and second-lowest total KLD variance. B) Heatmap of the RF importance for all features of original data (reproduced from Fig 1F) and all synthetic data-generating approaches.

Figure S6C displays the result of projecting synthetic-feature principal components onto the feature principal components of the original data. Here, a synthetic technique that leads to distinct clustering from the original PCA analysis suggests that the features do not preserve the local structure. We find that all synthetic datasets produce nearly overlapping ellipses by this PCA analysis. Individual PCA analyses performed independently on each synthetic matrix are shown in Figure S8. Comparing the PCA analyses in Figure 1C, we find the supervised synthetic approaches SMOTE and KNN (with k = 20) reproduce the PCA subclustering pattern. This presents an interesting juxtaposition between two different methods of evaluating synthetic data: one based on a quantitative measure (KLD) and another based on a visual/qualitative assessment (PCA plot comparison). The KLD analysis suggests that GMM is capturing the overall statistical properties of the data the best (SMOTE being the worst), while the PCA analyses suggests data generated by SMOTE and KNN (with k = 20) better captures the cluster structure when the data is projected onto the first two principal components.

Figure 4B shows the importance measures of all synthetic datasets used in the RF models, with the RF model training performed similarly as in the previous sections (see Methods). We find that GMM and MVN lead to overemphasized importance for visit 4 saliva RBD IgG, visit 4 serum IgG RBD, and a visit 1 serum IgG RBD. SMOTE appears to perform overall quite well in terms of reproducing the importance feature distribution, with all RF feature importance metrics comparing best to the original data.

## 2 Discussion

Our implementation of random forest (RF) is capable of accurately classifying the SARS-CoV-2 vaccination response of PLWH from an age-matched HIV-negative group with humoral and cellular immunological data from SARS-CoV-2 vaccination. Our ablation procedures, based on feature importance, reveal an optimized subset of features comprising IL2 and IFNg cytokine-producing T cells, the CD8/CD4 ratio, and saliva IgA RBD, that lead to equivalent RF model performance as the full 63 feature dataset (Figure 3A-C). Thus, cytokine-based features in combination with post-booster saliva IgA features, contain most of the predictive power to differentiate PLWH immunogenic responses in our dataset. Adding additional features beyond the optimal 9 appears to have little effect on the AUC, but causes the F1 score distribution to deviate from the full-feature model F1 Score distribution until the 35th feature is added. We believe this to be a commonly observed phenomenon in which additional features beyond the optimal subset confuse the model and degrade performance [26]. Thus, the ablation procedure finds the 9 optimal features contain most of the classification power, with the remaining 54 features contributing mostly minimal or redundant information. The eventual improvement at 35 features (Figure 3B, blue squares) indicates that additional useful information is present but only becomes beneficial when enough features are considered together, suggesting some further interactions or combinations of immunological responses are relevant.

Analyzing individual classification probabilities reveals there is a select group of individuals that the RF algorithm has difficulty correctly classifying (IDs 39, 72, 6 from Figure 3C). Consistently misclassified PLWH may reflect atypical immune responses, such as elite controllers or unique treatment effects, while misclassified HIV-negative individuals may have immune profiles resembling HIV-positive cases due to comorbidities like chronic inflammation or autoimmune conditions; therefore, the RF algorithm can be used to identify these misclassified individuals for closer clinical follow-up to uncover unique immune mechanisms, guide personalized care, and address potential comorbidities or atypical disease progression. This optimization framework lays the foundation for immune system monitoring in longitudinal vaccine studies when balancing comprehensive immunological profiling in resource-limited settings. The resulting sparse model not only preserves the predictive power of the RF classifier but also delivers a feasible approach for vaccine response when the high costs of clinical studies are challenging. As such, identifying key predictive parameters fosters robust immunological surveillance by simultaneously shrinking model complexity and sampling burden.

We observed that immunological feature types that form distinct clusters by t-SNE, such as cytokines, saliva IgG, and saliva IgA (Figure 2A) also appear to produce relatively high AUC-ROC’s when training RF models in isolation (Figure 3D). In contrast, clinical features that do not form distinct clusters by t-SNE, such as serum IgG and neutralization/ACE2 displacement, lead to relatively poor median AUC-ROC performance (Figure 3D). Therefore, there may be an underlying relationship between the high dimensional t-SNE feature clustering and RF predictive power. t-SNE is an unsupervised method used to visually capture the underlying local and global structure inherent to a dataset [27]. RF models tend to perform optimally when provided with informative non-redundant features [28]. t-SNE, by revealing local cluster formation, may be implicitly revealing subsets of features that have low internal noise and high predictive relevance. Indeed, previous work has successfully implemented an initial t-SNE screening step to eliminate redundant data from a large reference dataset prior to RF model training [29]. Supervised and unsupervised clustering may therefore warrant further exploration in immunological context to reduce redundancy in datasets prior to model training.

T-cell responses play a crucial role in shaping immunity after mRNA COVID-19 vaccination [30], with spike-specific IFNg and IL2 T-cell responses emerging as key factors following repeated vaccination [31, 32]. In this work, we use ELISpot cytokine responses to SARS-CoV-2 spike peptides as a measure of frequency of spike-specific T cells. We found the most important features to distinguish PLWH vaccine-specific immunological outcomes in our RF approach to be the post-primary series and booster IL2 responses and post-booster IFNg responses appear to carry the strongest classifying power resulting in a median AUC-ROC of ∼0.97 and F1 Score of ∼0.8 (Figure 3A). Cytokine-based statistical differences between PLWH and the HIV-negative individuals were previously documented with this dataset and found to deviate significantly from the HIV-negative individuals, with the underlying mechanism hypothesized to be Th1 imprinting from pre-existing HIV infection [7]. Clinicians are more often utilizing ROC curves to make diagnostic judgements [33]. Our ROC (Figure 3E) and PR (Figure S5) curves computed from RF-training on cytokine features indeed show the cytokine features to be diagnostically near-perfect classifiers in the context of vaccine-elicited immunity among PLWH.

Previous work on immunogenic outcomes from SARS-CoV-2 vaccination reveals that PLWH mounts overall similar serum IgG immunity to vaccination [7, 19–22], with serum IgG production and decay rates estimated via mechanistic modelling also found to be similar to HIV-negative individuals [7]. Where serum IgG spike and RBD features are the most populated feature type in our work (28 features in total), all of them are found to be the least-informative features for RF model classification (Figure 1F), with no median weights deviating from ∼0 during training (Figure 2D,E). We therefore find results from the RF approach to be in agreement with previous literature for serum-based IgG features with PLWH mounting indistinguishable IgG responses from an age-matched HIV-negative group. Serum IgG is primarily produced by B cells in the bone marrow, spleen, and lymph nodes [34, 35]. Despite evidence for T-cell dysregulations due to ART-suppressed HIV viremia, there is likely effective restoration of the systemic immunity of these central immune organs associated with mounting serum IgG responses to a degree where vaccine-elicited serum IgG responses are non-informative via a RF approach.

Gut-associated lymphoid tissue contains a distinct repertoire of lymphocytes that regulate mucosal humoral immunity [36], further, plasmablasts responsible for IgA secretion are believed to migrate between bone marrow, blood, and mucus [37]. IgA is the first line of defense against SARS-CoV-2 infection [38, 39]. During the initial stages of HIV-1 infection, CD4+ T cell in the mucusal immune system have been shown to be more significantly impacted than systemic CD4+ T cells [40], further, IgA production and maintenance is known to be dysregulated by HIV-1 [41]. The extent to which mucosal immunity is restored, or further dysregulated, among PLWH on ART is not fully known [42, 43]. In the context of live-attenuated influenza vaccines in healthy young adults, mucosal and systemic humoral responses were found to be regulated by separate and distinct mechanisms [44]. We were therefore interested in determining if a RF approach reveals a unique saliva-based immunogonenic signature among HIV+ ART-suppressed individuals who received repeated SARS-CoV-2 vaccinations. In the dataset used in this work, the overall amplitude of saliva IgA responses both pre and post primary series remained relatively unchanged and was not determined to be a strong statistical differentiator between PLWH and the control group, whereas some significant statistical differences for a subset of IgG visits were reported [7]. However, our ablation analysis revealed saliva IgA measures to contribute to the minimally stratified combination of features, suggesting saliva IgA are highly informative towards differentiating PLWH vaccine immunologic response from the HIV-negative control. This result may be partially explained by previous clinical literature. For example, PLWH exhibit chronic antigenic stimulation leading to B-cell dysregulation and reduced IgA-producing plasmablasts [45], where the proposed mechanism effecting IgA production is HIV nef protein penetrating B cells and blocking cytokine signalling [46]. Thus, our findings that saliva-based IgA is a key biomarker to distinguish a PLWH immunogenicity highlights the utility of RF to detect subtle dysregulations in mucosal immune profiles not previously found by statistical and mechanistic modelling approaches [7].

ML-generated synthetic data have incredible promise to increase diversity and robustness in medical and healthcare datasets, as well as make reproducibility of results more achievable through datasets that can be shared while preserving the privacy of protected health information [47, 48]. The human immune system is incredibly complex, and the degree to which its myriad of components are functionally related is not well known [49]. ML techniques that can capture these underlying relationships are valuable for overcoming data limitations for training, model validation, and reducing ethical constraints on data sharing. Furthermore, mechanistic modelling of the relationship between clinical outcomes and underlying immunological processes is often limited due to practical identifiability from data scarcity [50], particularly in longitudinal vaccine studies where frequent sampling and comprehensive immune profiling may not be feasible [32, 51–56]. Therefore, synthetic data that preserves the local and global characteristics of the original dataset’s structure would enable *in silico* testing of immunological hypotheses through computational approaches. We explore synthetic data generation using unsupervised methods such as GMM, MVN, and KNN, as well as supervised methods such as SMOTE. Surprisingly, we find that SMOTE produces the least favorable outcome as quantified by KLD divergence, but the most favorable outcome by qualitative PCA comparison to the original data (Figure S8C versus Figure 1C) as well as quantitative comparison of feature importance behaviour from RF model outcomes (Figure 4B). GMM achieves the lowest KLD (Figure S8A), however, it fails to preserve the data structure from PCA analyses (Figure S6A) and leads to inflated feature importance scores (Figure 4B). Our results underscore how different approaches may capture various aspects of the global and local structure within the data and further show that different synthetic data generation approaches capture distinct aspects of the immunological response. A hybrid approach-combining the strength of GMM in modeling global distributions with SMOTE’s ability to preserve local feature relationships - might better recapitulate the complex immune signatures observed in vaccine responses. Future work will explore such hybrid methodologies alongside advanced deep learning approaches, such as variational autoencoders (VAEs), generative adversarial networks (GANs), diffusion and energy-based models, to further enhance generative capabilities and improve model robustness.

In this paper, we demonstrate RFs are able to identify key immunogenic differences between PLWH and an age-matched HIV-negative group. ML techniques have great potential to pinpoint critical biomarkers through classification and clustering approaches, furthering the depth of our understanding of immunogenic longitudinal outcomes. ML approaches hold significant promise in the field of immunology due to their ability to uncover complex patterns in large, high-dimensional datasets that may not be readily apparent through mechanistic modelling approaches [57–60]. As novel vaccine therapeutics are optimized and innovated [61], ML approaches can be utilized to better understand vaccine-elicited immune responses.

## 3 Methods

### 3.0.1 Study Approval

Vaccinations were not provided as a part of this study. All study participants provided informed written consent. The study protocol and consent form were approved by the University of Toronto Research Ethics Board (RIS #40713) and Sinai Health REB (21-0223-E).The study was conducted in accordance with the protocol, applicable regulations, and guidelines for Good Clinical Practice (GCP), Health Canada’s regulations, and the Tri-Council Policy Statement: Ethical Conduct for Research Involving Humans (TCPS 2.0).

### 3.1 Clinical data acquisition and description

This study uses SARS-CoV-2 vaccination data for three doses of vaccine previously published in ref. [7]. Primary booster series data, corresponding to Visits 1 through 9 were acquired for this work through a data sharing agreement between authors JMH and MO. Visit acquisition times can be found in the timeline schematic in Figure 1A. Briefly, visits 1-9 occur over 48 weeks post dose 1 vaccination, and consist of individuals who received 3 doses of SARS-CoV-2 vaccination. These data comprise 91 individuals: 23 HIV- and 63 HIV+. Individuals are predominantly male, recruited from a clinic with a large MSM population. Visits 10-11 IgG spike and RBD serum data corresponding to post-SARS-CoV-2 vaccinations 4 and 5, respectively, were also used in this work. These data were collected by the A.C.G. lab in a follow up study to ref. [7] (see acknowledgements) and are unpublished primary data. The timeline of study protocol can be found in Figure 1B.

#### 3.1.1 Antibody detection in serum

Antibody detection in serum for visits 10-11 were carried out as described in ref. [7]. An automated ELISA assay was employed to measure total IgG antibody levels against the full-length spike trimer, RBD, and nucleocapsid, as described previously [62, 63] In summary, 384-well microplates were precoated with spike (SmT1), RBD, or nucleocapsid antigens provided by the National Research Council of Canada (NRC). Key steps included blocking with Blocker BLOTTO (ThermoFisher Scientific), incubation with serum dilutions of 1:160 or 1:2,560, followed by incubation with HRP-conjugated human anti-IgG (IgG#5, supplied by NRC). Detection was carried out using ELISA Pico Chemiluminescent Substrate (ThermoFisher Scientific), and chemiluminescence was measured with an EnVision 2105 Multimode Plate Reader (Perkin Elmer). Raw chemiluminescence values were normalized using a synthetic standard included on each plate (VHH72-Fc from NRC for spike/RBD or anti-N IgG from Genscript, #A02039). These values were further converted to BAU/mL using the WHO International Standard 20/136 as a calibrator [62]. Seropositivity thresholds for the 1:160 dilution were established based on 3 standard deviations (3SD) above the mean of control samples [62].

### 3.2 Missing data imputation

Clinical features with missing entries are common when dealing with immunological clinical data [64] with the imputation necessary to carry out basic ML algorithms [65]. We impute missing data using the **R** library missForest [66] (version 1.5) to each class (HIV- and HIV+) separately. missForest works by iteratively training a Random Forest model on the observed data to predict and replace missing values for each feature, cycling through all features until convergence. Imputation of clinical data using missForest has been widely implemented and shown to handle mix-type data well [67–69]. In this work, all 63 of our clinical features are continuous variables. Normalized root mean squared error (NRMSE) was computed to assess imputation accuracy. The NRMSE for the HIV- and HIV+ imputed values were calculated to be 0.38 and 0.36, respectively. The most commonly imputed feature types were the saliva antibody features. In order to determine the accuracy of imputed values, in the supplementary material we assess the influence of low frequency of mutual missing values on imputed results through randomly removing observed values from all feature compartments and impute them using missForest to measure the model accuracy. The NRMSE falls within a reasonable range for missForest’s performance as a function of increasing percentage of missingness (Fig S9), and the mean and variance of all biomarkers remain stable across increasing levels of missingness (Fig S10 and S11, respectively).

### 3.3 Correlation network visualization

We visualize the correlation network structure of the clinical features, with each features represented as a node, and with the network layout calculated using the t-distributed Stochastic Neighbor Embedding (t-SNE) algorithm [27] applied to the adjacency matrix. It is recommended t-SNE perplexity satisfy 3 ∗ perplexity *<* feature count − 1 [70], we therefore set the perplexity to (63 − 1)*/*3 where 63 is the number of features. The network node size is proportional to RF feature importance. Vertex colour-code varies and is described in the respective figure caption. For visualization purposes, we only plot an edge between a pair of features if the correlation is statistically significant (p-values *<* 0.05). The t-SNE layout is implemented using the *Rtsne* function from the Rtsne library in **R**.

### 3.4 Model development and model performance metrics

Principal component analysis (PCA) is an unsupervised dimensionality reduction technique that transforms high-dimensional data into orthogonal components that preserves as much variance as possible. We perform PCA on the full dataset to validate whether an unsupervised clustering technique leads to separate clusters composed of our data classes (HIV+ and HIV-). PCA is implemented using the **R** function *prcomp* in the stats library, with confidence ellipses of levels of one standard deviation drawn using the *dataEllipse* function in the car library. Linear discriminant analysis (LDA) is used to provide initial insights into class separability. LDA is implemented using the *lda* function in the MASS library in **R**.

We employed the RF algorithm [71]; a nonlinear ensemble-based method, to quantify the complex nonlinear relationships between HIV+ and HIV-immunological responses. All training sets are randomly downsampled to balanced population sizes of 20:20 for HIV-:HIV+ individuals. The number of trees was initially increased until the ORR error stabilized and was then fixed throughout this work to a value of 200. Random holdouts of population proportions of 3:12 (HIV-:HIV+) were excluded from training and used to assess model performance. 1000 stochastically generated instances of the RF algorithm (each with randomly selected training and testing sets) was considered to assess RF model performance. Majority tree visualization was explored to ‘look under the hood’ of every given RF-training and RF-testing landscapes to assess how features were being weighted for each randomly sampled training and and holdout (test) set. To do this, we construct a model explanation function using the *lime* and *explain* functions from the lime library in **R**. Feature weights from every RF model were then stored and visualized (see supplementary for individual model examples). For all RF models, a threshold of 0.5 is used. The repercussion of a 0.5 threshold on individual prediction accuracy with a 0.5 threshold are explored in the supplementary material.

An underlying assumption of the RF algorithm is the statistical independence of all observations. In this study, while the patients are independent, the samples collected from various vaccines for the same patient are not. To address this, we ensure that samples from the same patient taken during multiple vaccine trials remain within the same fold to prevent them from being dispersed across both training and test datasets. Furthermore, we implemented a two-layer cross-validation procedure designed to optimize model hyperparameters while ensuring that predictions are made exclusively on samples not seen during the training process to guarantee complete independence from intra-patient correlations.

### 3.5 Feature importance

Gini Impurity measures the likelihood of incorrect classification of a randomly chosen element if it were labeled according to the class distribution in a dataset. RF splits the dataset at each node using a feature that minimizes Gini Impurity. At each split, the Gini Impurity before and after the split is calculated, and the reduction in Gini Impurity due to that split is recorded. Feature importance in RF is derived by summing the total reduction in Gini Impurity across all trees in the forest, weighted by the number of samples passing through each node. This was implemented using the *varImp* function in **R**. The cumulative average from each importance calculation was then calculated across all RF models.

### 3.6 Ablation analysis

To assess sensitivity of RF performance on our features, and further, to identify and select the most important features that contribute the most to the model accuracy and performance, we implemented two types of ablation analysis. Reverse ablation, whereby features are removed as a function of most importance until the two least important features remain, was conducted to determine the minimal accuracy of our 2 least important features. Forward ablation, which begins with the two most important features then iteratively adds in features as a function of their importance, was conducted to determine the subset of features that lead to equivalent model performance.

For every ablation index, the RF model is employed as described above with 1000 stochastically generated instances of training and test sets. Equivalent model performance was assessed by computing a Wilcoxon Test between the resulting distributions of the AUC-ROC and F1 Scores and their respective ‘full dataset’ counter parts for the forward ablation algorithm. P-values below 0.05 thus indicate non-equivalent model performance, where p-values above 0.05 indicate equivalent model performance to the full 63 feature dataset. Where p-values cross from below to above 0.05 therefore suggests the minimal subset of features to achieve equivalent model performance when using the full 63 feature dataset.

### 3.7 Simulated data

A major motivation for this study was to develop a robust framework for generating synthetic immunological data that assimilates the complex patterns observed in our original vaccine response dataset. We implemented 6 synthetic data generating techniques: Multivariate Normal (MVN), Gaussian Mixture Model (GMM), Synthetic Minority Oversampling Technique (SMOTE), and K-nearest neighbours (KNN) with *k* = 3, *k* = 10 (determined as the optimal *k* value), and *k* = 20 (2x optimal). For unsupervised techniques, HIV- and HIV+ data were generated separately from one another. All synthetic datasets are generated equal to our original dataset’s class balance and feature size: each are implemented to uniquely generate 91 individuals (23 HIV- and 68 HIV+) as well as 63 corresponding immunological features. SMOTE is a supervised technique typically used to generate synthetic data to achieve a balanced dataset. Here, we apply SMOTE to generate synthetic samples for both the minority and majority classes. MVN, GMM, SMOTE, and KNN are all implemented in **R** using the `MASS`, `mclust`, `smotefamily`, and `FNN` libraries, respectively.

### 3.8 Evaluation of simulated data approaches

We use Kullback–Leibler divergence (KLD) to evaluate and rank the synthetic data methods by measuring how much the original data is different from the synthetic dataset that we have generated via the various methods described above. The KLD is calculated using the *KLD* function in the `LaplacesDemon` library in **R**. We calculate the KLD between all 63 synthetic and original features. The mean KLD is then computed by taking the mean of all the KLD for each synthetic approach, similarly, we also compute the total variance across all KLD for each synthetic approach. Synthetic approaches are then ranked by their mean and total variance KLDs: a low mean KLD, computed across all features, suggests that the synthetic data approach is close to the original distribution of features, while a low total KLD variance means that the KLD values across features being synthetically produced contain a consistent level of similarity to the original data. Low total KLD variance also suggests that there are fewer subsets of features deviating significantly from the original data. Finally, for the best-ranked synthetic method we compute variance in KLD as a function of feature type to assess if certain synthetic features are closer to the original data than others.

We also utilize PCA to assess how similar synthetic data features are to the original data. To do this, we perform PCA on the original data and then project each synthetic dataset onto the PCA space of the original dataset. Ellipses drawn to the first standard deviations of the resulting first two principal components are then used to visually inspect clustering behaviour. If the ellipses representing the synthetic data generated by one method overlap significantly with the ellipses from the original data in the PCA space, it suggests that the synthetic data closely resembles the distribution and structure of the original data in the reduced dimensionality space.

## 4 Data and Code Availability

The data and code used to produce this work will be made publicly available, please stay tuned to updates to this manuscript for an active link to the public Github repository. Please email the corresponding author if you want access as soon as it is available.

### 5 Supplementary information

Supplementary information and supporting text accompanies this manuscript.

### 6 Author Contributions

Conceptualization: C.S.K., J.C., M.S.G., J.M.H. Funding Acquisition: C.S.K., M.O., J.M.H. Data Curation: C.S.K. & V.M. Visualization: C.S.K. & M.S.G. Methodology: C.S.K., J.C., J.M.H., M.S.G. Investigation: C.S.K., M.S.G., J.C., J.M.H., V.M., M.O. Validation: C.S.K., V.A.M. Formal Analysis: C.S.K., M.S.G. Supervision: M.S.G., J.M.H., & J.C. Writing – Original Draft Preparation: C.S.K. Writing - Review & Editing: All authors.

## 8 Acknowledgments

This project is supported by the NRC-Fields Mathematical Sciences Collaboration Centre and the AI4PH program, sponsored by the Dalla Lana School of Public Health, University of Toronto. C.S.K. acknowledges NSERC postdoctoral funding. J.M.H. acknowledges funding from NSERC, CIHR, NSERC EIDM, and the York Research Chair Program. J.M.C. acknowledges the support of the National Institutes of Health (grant nos. R21-AI143443-01A1 and R01-OD011095). The authors are thankful to the Anne-Claude Gingras group (Lunenfeld-Tanenbaum Research Institute, Sinai Health, Toronto, Canada) for the IgG ELISA on sera from the two final study visits.

## 9 Materials & Correspondence

## References

[1] Bekker, L.-G., Beyrer, C., Mgodi, N., Lewin, S.R., Delany-Moretlwe, S., Taiwo, B., Masters, M.C., Lazarus, J.V.: HIV infection. Nature Reviews Disease Primers 9(1), 42 (2023). 10.1038/s41572-023-00452-3

[2] Carter, A., Zhang, M., Tram, K.H., Walters, M.K., Jahagirdar, D., Brewer, E.D., Novotney, A., Lasher, D., Mpolya, E.A., Vongpradith, A., Ma, J., Verma, M., Frank, T.D., He, J., Byrne, S., Lin, C., Dominguez, R.-M.V., Pease, S.A., Comfort, H., May, E.A., Abate, Y.H., Abbastabar, H., Abdelkader, A., Abdi, P., Abdoun, M., Abdul Aziz, J.M., Abidi, H., Abiodun, O., Aboagye, R.G., Abreu, L.G., Abtew, Y.D., Abu-Gharbieh, E., Aburuz, S., Abu-Zaid, A., Addo, I.Y., Adegboye, O.A., Adekanmbi, V., Adetunji, C.O., Adetunji, J.B., Adeyinka, D.A., Adhikari, K., Adnani, Q.E.S., Adzigbli, L.A., Afrashteh, F., Afzal, S., Aghamiri, S., Agide, F.D., Agodi, A., Agyemang-Duah, W., Ahinkorah, B.O., Ahmad, F., Ahmad, S., Ahmad, S., Ahmad, A., Ahmed, I., Ahmed, H., Ahmed, S.A., Ahmed, S., Ahmed, A., Ahmed, M., Ahmed, A., Akalu, G.T., Akinosoglou, K., Al Awaidy, S., Al Hamad, H., Al Mosa, A.S., Al Zaabi, O.A.M., Alalalmeh, S.O., Alam, N., Alam, N., Alanezi, F.M., Alayu, D.S., AlBataineh, M.T., Alemohammad, S.Y., Al-Gheethi, A.A.S., Ali, S.S., Ali, M.U., Ali, A., Ali, L., Ali, W., Al-Ibraheem, A., Almazan, J.U., Altaf, A., Altwalbeh, D., Alvis-Guzman, N., Al-Zyoud, W.A., Amani, R., Amera, T.G., Ameyaw, E.K., Amiri, S., Amu, H., Amusa, G.A., Anil, A., Anjorin, A.-A.A., Antonio, C.A.T., Anwar, S., Anwer, R., Anyabolo, E.E., Anyasodor, A.E., Apostol, G.L.C., Ardekani, A., er Areda, Aregawi, B.B., Aremu, A., Armani, K., Asemahagn, M.A., Ashemo, M.Y., Ashraf, T., Asika, M.O., Asmerom, H.A., Atout, M.M.W., Aujayeb, A., Awad, H., Awotidebe, A.W., Ayala Quintanilla, B.P., Ayele, F., Azadnajafabad, S., Aziz, S., B, D.B., Babu, G.R., Badar, M., Bahramian, S., Bako, A.T., Balcha, W.F., Bam, K., Banik, B., Bardhan, M., Bärnighausen, T.W., Barqawi, H.J., Basharat, Z., Bashiru, H.A., Basiru, A., Bastan, M.-M., Basu, S., Bathini, P.P., Batra, K., Batra, R., Bayleyegn, N.S., Begum, T., Behnoush, A.H., Beiranvand, M., Belete, M.A., Belete, A.C., Beloukas, A., Beneke, A.A., Beran, A., Berhie, A.Y., Bermudez, A.N.C., Bernstein, R.S., Beyene, K.A., Bhardwaj, P., Bhardwaj, N., Bhat, A.N., Bhat, V., Bhatti, G.K., Bhatti, J.S.S., Bishaw, K.A., Bisht, K.D., Bodhare, T., Bodunrin, A.O., Boltaev, A.A., Borhany, H., Bouaoud, S., Brown, C.S., Buonsenso, D., Burkart, K., Bustanji, Y., Butt, Z.A., Cao, C., Cárdenas, R., Cenderadewi, M., Chadwick, J., Chakraborty, C., Chakraborty, S., Chandika, R.M., Chattu, V.K., Chaurasia, A., Chen, G., Ching, P.R., Chopra, H., Choudhari, S.G., Chu, D.-T., Chukwu, I.S., Chung, E., Cindi, Z., Couto, R.A.S., Cruz-Martins, N., Cuadra-Hernández, S.M., Dabo, B., Dadras, O., Dagnew, G.W., Dahiru, T., Dai, X., Darwesh, A.M., das Neves, J., Dash, N.R., Dashti, M., De la Hoz, F.P., Debopadhaya, S., Degenhardt, L., Delgado-Enciso, I., Deribe, K., Des Jarlais, D.C., Desai, H.D., Deuba, K., Dhane, A.S., Dhingra, S., Diaz, D., Diaz, M.R., Ding, D.D., Do, T.C., Dohare, S., Dongarwar, D., dos Santos, W.M., Doshi, O.P., Dsouza, A.C., Dsouza, H.L., Dsouza, V.S., Duraisamy, S., Dziedzic, A.M., Ebrahimi, A., Ed-Dra, A., Edinur, H.A., Efendi, F., Ekholuenetale, M., Ekundayo, T.C., El Sayed, I., Elhadi, M., Eltaha, C., Eskandarieh, S., Eslami, M., Eze, U.A., Fahim, A., Fatehizadeh, A., Fauk, N.K., Fazeli, P., Fekadu, G., Ferreira, N., Firew, B.S., Fischer, F., Folayan, M.O., Foroutan, B., Fukumoto, T., G, S., Gadanya, M.A., Gaidhane, A.M., Gaipov, A., Gandhi, A.P., Ganiyani, M.A., Gebregergis, M.W., Gebrehiwot, M., Gebremeskel, T.G., Getachew, M.E., Ghadiri, K., Ghasemzadeh, A., Ghashghaee, A., Gholami, E., Gholizadeh, N., Ghorbani, M., Gil, A.U., Girmay, A.A., Golechha, M., Golinelli, D., Goulart, A.C., Goyal, A., Gudeta, M.D., Gupta, S., Gupta, B., Habteyohannes, A.D., Haghmorad, D., Haj-Mirzaian, A., Halwani, R., Handiso, D.W., Haq, Z.A., Harapan, H., Hargono, A., Hasaballah, A.I., Hasnain, M.S., Hassan, S., Hassanipour, S., Hegazi, O.E., Heidari, M., Hezam, K., Hlongwa, M.M., Hoan, N.Q., Hoogar, P., Hosseinzadeh, M., Hosseinzadeh Adli, A., Hundie, T.G., Hushmandi, K., Huynh, H.-H., Ibitoye, S.E., Ikiroma, A., Ikuta, K.S., Ilesanmi, O.S., Ilic, I.M., Iradukunda, A., Isa, M.A., Ismail, N.E., Iyamu, I.O., J, V., Jacobsen, K.H., Jain, A., Jairoun, A.A., Jakovljevic, M., Janodia, M.D., Javadi Mamaghani, A., Jema, A.T., Jokar, M., Jonas, J.B., Joseph, N., Joshua, C.E., Kabir, A., Kabir, M.A., Kabir, Z., Kadashetti, V., Kaliyadan, F., Kanmodi, K.K., Kannan S, S., Karaye, I.M., Karimi Behnagh, A., Kassel, M.B., Kayode, G.A., Khajuria, H., Khalid, N., Khalil, A.A., Khamesipour, F., Khan, G., Khan, E.A., Khan, Y.H., Khan, M.J., Khan, M.N., Khatab, K., Khidri, F.F., Khorrami, Z., Khosravi, M., Khubchandani, J., Kim, M.S., Kim, J.Y., Kim, Y.J., Kisa, A., Kisa, S., Komaki, S., Kondlahalli, S.K.M.M., Koul, P.A., Koulmane Laxminarayana, S.L., Krishan, K., Kuate Defo, B., Kuddus, M.A., Kulimbet, M., Kulkarni, V., Kumar, R., Kumar, V., Kumar, N., Kumar, M., Ladan, M.A., Lal, D.K., Le, T.T.T., Le, N.H.H., Lee, S.W., LeGrand, K.E., Lerango, T.L., Li, M.-C., Ligade, V.S., Lim, S.S., Limenh, L.W., Liu, X., Liu, R., Lodha, R., Loreche, A.M., M. Amin, H.I., Ma, Z.F., Majeed, A., Malakan Rad, E., Malhotra, H.S., Malhotra, K., Malik, A.A., Malik, I., Mallhi, T.H., Mansournia, M.A., Marasini, B.P., Martinez-Guerra, B.A., Martins-Melo, F.R.R., Martorell, M., Marzo, R.R., Mathur, N., McKowen, A.L.W., Meles, H.N., Melese, E.B., Memish, Z.A., Mendoza, W., Menezes, R.G., Meretoja, T.J., Mestrovic, T., Meylakhs, P., Mhlanga, L., Michalek, I.M., Micheletti Gomide Nogueira de Sá, A.C., Minervini, G., Minh, L.H.N., Moazen, B., Mohamed, N.S., MohammadAlizadeh-Charandabi, S., Mohammadian-Hafshejani, A., Mohammed, H., Mohammed, S., Mohammed, M., Mokdad, A.H., Monasta, L., Moni, M.A., Montazeri, F., Moradi, M., Moradi, Y., Motappa, R., Mougin, V., Mubarik, S., Mukoro, G.D., Mulita, F., Munjal, K., Munkhsaikhan, Y., Murlimanju, B.V., Musaigwa, F., Mustafa, G., Muthupandian, S., Nagarajan, A.J., Naghavi, P., Naik, G., Nainu, F., Najafi, M.S., Nargus, S., Navaratna, S.N.K., Naveed, M., Nayak, V.C., Nayak, B.P., Nduaguba, S.O., Negesse, C.T., Nematollahi, M.H., Nguefack-Tsague, G., Nguyen, D.H., Nguyen, H.Q., Nguyen, V.T., Niazi, R.K., Nigatu, Y.T., Nikravangolsefid, N., Niranjan, V., Nnaji, C.A., Noor, S.T.A., Not applicable, N., Noubiap, J.J., Nri-Ezedi, C.A., Nugen, F., Nutor, J.J., Nzoputam, C.I., Nzoputam, O.J., Obamiro, K.O., Odetokun, I.A., Oghenetega, O.B., Oguntade, A.S., Okeke, S.R., Okekunle, A.P., Okonji, O.C., Olagunju, A.T., Olakunde, B.O., Olalusi, O.V., Olatubi, M.I., Olorukooba, A.A., Olufadewa, I.I., Omar Bali, A., Onwujekwe, O.E., Opejin, A., Ordak, M., Orish, V.N., Ortiz-Brizuela, E., Osuagwu, U.L., Ouyahia, A., P A, M.P., Padubidri, J.R., Palladino, C., Pandey, A., Panos, L.D., Paredes, J.L., Parija, P.P., Parikh, R.R., Pashaei, A., Pasovic, M., Patel, S.K., Pathan, A.R., Patil, S., Pawar, S., Pepito, V.C.F., Peprah, E.K., Peprah, P., Pereira, M., Perna, S., Petcu, I.-R., Pham, H.T., Pillay, J.D., Poluru, R., Postma, M.J., Pourtaheri, N., Pradhan, J., Prakash, P., Prakasham, T.N.N., Prates, E.J.S., Pribadi, D.R.A., Priscilla, T., Puvvula, J., Qattea, I., Qazi, A.S., Radhakrishnan, R.A., Rafferty, Q., Rafique, I., Rahim, F., Rahimi-Movaghar, A., Rahimi-Movaghar, V., Rahman, M., Rahmani, A.M., Rahmani, S., Rahmanian, N., Rahmanian, M., Rahmanian, V., Rajaa, S., Ramadan, M.M., Ramadan, H., Ramasamy, S.K., Ramesh, P.S., Rana, K., Ranabhat, C.L., Rao, M., Rao, S.J., Rashidi, M.-M., Rathish, D., Rauniyar, S.K., Rawaf, S., Redwan, E.M.M., Reiner Jr., R.C., Rezaeian, M., Rodriguez, J.A.B., Root, K.T., Ross, A.G., Rotimi, K., Roy, N., Rwegerera, G.M., Sabet, C.J., Saddik, B.A., Saeb, M.R., Saeed, U., Saeedi, P., Safi, S.Z.Z., Sagar, R., Saheb Sharif-Askari, F., Saheb Sharif-Askari, N., Sahebkar, A., Sahoo, S.S., Saif, Z., Sajid, M.R., Salam, N., Salami, A.A., Saleh, M.A., Salehi, L., Samadi Kafil, H., Samy, A.M., Sanjeev, R.K., Santric-Milicevic, M.M., Saravanan, A., Sartorius, B., Sathyanarayan, A., Satpathy, M., Sawhney, M., Sedighi, M., Semagn, B.E., Senapati, S., Sethi, Y., Seylani, A., Shah, P.A., Shahid, S., Shaikh, M.A., Shamekh, A., Shamshirgaran, M.A., Shamsi, A., Shanawaz, M., Shannawaz, M., Sharifan, A., Sharifi-Rad, J., Shastry, S., Shenoy, R.R., Shetty, P.K., Shetty, M., Shetty, P.H., Shiferaw, D., Shirkoohi, R., Shittu, A., Shrestha, S., Sibhat, M.M., Siddig, E.E., Siedner, M.J., Singh, J.A., Singh, P., Singh, S., Singh, H., Sinto, R., Skryabina, A.A., Smith, A.E., Sobia, F., Sokhan, A., Solanki, S., Solanki, R., Sorensen, R.J.D., Sulaiman, S.K., Szarpak, L., T Y, S.S., Tabish, M., Tadakamadla, S.K., Taheri Abkenar, Y., Taiba, J., Talaat, I.M., Tampa, M., Tamuzi, J.L., Tan, K.-K., Tanwar, M., Tarkang, E.E., Taveira, N., Teklay, G., Tesfaye, B.T., Teye-Kwadjo, E., Thakur, R., Thangaraju, P., Thapa, R., Thapar, R., Thienemann, F., Thomas, J., Tovani-Palone, M.R., Tran, T.H., Tran, M.T.N., Tsai, A.C., Tsegay, G.M., Tumurkhuu, M., Udoh, A., Ullah, I., Ullah, A., Umair, M., Umar, M., Unnikrishnan, B., Vahdati, S., Vaithinathan, A.G., Varthya, S.B., Vasankari, T.J., Verras, G.-I., Villafañe, J.H., Vo, A.T., Vos, T., Walde, M.T., Wamai, R.G., Wang, Y., Waqas, M., Ward, P., Wassie, G.T., Weintraub, R.G., Weldetinsaa, H.L., Weldu, G.A., Westerman, R., Wickramasinghe, N.D., Woldekidan, M.A., Wong, Y.J., Worku, N.K., Wu, Z., Wu, X., Yaghoubi, S., Yesera, G.E., Yezli, S., Yi, S., Yiğit, A., Yin, D., Yismaw, Y., Yon, D.K., Yonemoto, N., Zakham, F., Zhang, H., Zhang, J., Zhao, H., Zhu, B., Zhuang, Q., Zhumagaliuly, A., Zielińska, M., Zihao, L., Zikarg, Y.T., Zoladl, M., Zumla, A., Zyoud, S.H., Zheng, P., Aravkin, A.Y., Imai-Eaton, J.W., Naghavi, M., Schumacher, A.E., Hay, S.I., Murray, C.J.L., Kyu, H.: Global, regional, and national burden of HIV/AIDS, 1990&#x2013;2021, and forecasts to 2050, for 204 countries and territories: the Global Burden of Disease Study 2021. The Lancet HIV 11(12), 807–822 (2024). 10.1016/S2352-3018(24)00212-1

[3] Teeraananchai, S., Kerr, S.J., Amin, J., Ruxrungtham, K., Law, M.G.: Life expectancy of HIV-positive people after starting combination antiretroviral therapy: a meta-analysis. HIV medicine 18(4), 256–266 (2017). 10.1111/hiv.12421

[4] Deeks, S.G., Overbaugh, J., Phillips, A., Buchbinder, S.: HIV infection. Nature Reviews Disease Primers 1(October) (2015). 10.1038/nrdp.2015.35

[5] Bono, V., Augello, M., Tincati, C., Marchetti, G.: Failure of CD4+ T-cell Recovery upon Virally-Effective cART: an Enduring Gap in the Understanding of HIV+ Immunological non-Responders. New Microbiologica 45(3), 155–172 (2022)

[6] Lv, T., Cao, W., Li, T.: HIV-Related Immune Activation and Inflammation: Current Understanding and Strategies. Journal of Immunology Research 2021 (2021). 10.1155/2021/7316456

[7] Matveev, V.A., Mihelic, E.Z., Benko, E., Budylowski, P., Grocott, S., Lee, T., Korosec, C.S., Colwill, K., Stephenson, H., Law, R., Ward, L.A., Sheikh-Mohamed, S., Mailhot, G., Delgado-Brand, M., Pasculescu, A., Wang, J.H., Qi, F., Tursun, T., Kardava, L., Chau, S., Samaan, P., Imran, A., Copertino, D.C., Chao, G., Choi, Y., Reinhard, R.J., Kaul, R., Heffernan, J.M., Jones, R.B., Chun, T.-W., Moir, S., Singer, J., Gommerman, J., Gingras, A.-C., Kovacs, C., Ostrowski, M.: Immunogenicity of COVID-19 vaccines and their effect on HIV reservoir in older people with HIV. iScience 26(10), 107915 (2023). 10.1016/J.ISCI.2023.107915

[8] Saout, C.L., Lane, H.C., Catalfamo, M.: The role of cytokines in the pathogenesis and treatment of HIV infection. Cytokine and Growth Factor Reviews 23(4-5), 207–214 (2012). 10.1016/j.cytogfr.2012.05.007

[9] Kerńeis, S., Launay, O., Turbelin, C., Batteux, F., Hanslik, T., Böelle, P.Y.: Long-term immune responses to vaccination in HIV-infected patients: A systematic review and meta-analysis. Clinical Infectious Diseases 58(8), 1130–1139 (2014). 10.1093/cid/cit937

[10] El Chaer, F., El Sahly, H.M.: Vaccination in the Adult Patient Infected with HIV: A Review of Vaccine Efficacy and Immunogenicity. The American Journal of Medicine 132(4), 437–446 (2019). 10.1016/j.amjmed.2018.12.011

[11] Geretti, A.M., Doyle, T.: Immunization for HIV-positive individuals. Current Opinion in Infectious Diseases 23(1) (2010)

[12] García, F., Climent, N., Guardo, A.C., Gil, C., Léon, A., Autran, B., Lifson, J.D., Martínez-Picado, J., Dalmau, J., Clotet, B., Gatell, J.M., Plana, M., Gallart, T.: A dendritic cell-based vaccine elicits T cell responses associated with control of HIV-1 replication. Science Translational Medicine 5(166) (2013). 10.1126/scitranslmed.3004682

[13] Niessl, J., Baxter, A.E., Mendoza, P., Jankovic, M., Cohen, Y.Z., Butler, A.L., Lu, C.L., Dubé, M., Shimeliovich, I., Gruell, H., Klein, F., Caskey, M., Nussenzweig, M.C., Kaufmann, D.E.: Combination anti-HIV-1 antibody therapy is associated with increased virus-specific T cell immunity. Nature Medicine 26(2), 222–227 (2020). 10.1038/s41591-019-0747-1

[14] Höft, M.A., Burgers, W.A., Riou, C.: The immune response to SARS-CoV-2 in people with HIV. Cellular and Molecular Immunology 21(2), 184–196 (2024). 10.1038/s41423-023-01087-w

[15] Pollard, A.J., Bijker, E.M.: A guide to vaccinology: from basic principles to new developments. Nature Reviews Immunology 21(2), 83–100 (2021). 10.1038/s41577-020-00479-7

[16] Cox, R.J., Brokstad, K.A.: Not just antibodies: B cells and T cells mediate immunity to COVID-19. Nature Reviews Immunology 20(10), 581–582 (2020). 10.1038/s41577-020-00436-4

[17] Brodin, P., Davis, M.M.: Human immune system variation. Nature Reviews Immunology 17(1), 21–29 (2017). 10.1038/nri.2016.125

[18] Alirezaylavasani, A., Skeie, L.G., Egner, I.M., Chopra, A., Dahl, T.B., Prebensen, C., Vaage, J.T., Halvorsen, B., Lund-Johansen, F., Tonby, K., Reikvam, D.H., Stiksrud, B., Holter, J.C., Dyrhol-Riise, A.M., Munthe, L.A., Kared, H.: Vaccine responses and hybrid immunity in people living with HIV after SARS-CoV-2 breakthrough infections. npj Vaccines 9(1), 185 (2024). 10.1038/s41541-024-00972-3

[19] Frater, J., Ewer, K.J., Ogbe, A., Pace, M., Adele, S., Adland, E., Alagaratnam, J., Aley, P.K., Ali, M., Ansari, M.A., Bara, A., Bittaye, M., Broadhead, S., Brown, A., Brown, H., Cappuccini, F., Cooney, E., Dejnirattisai, W., Dold, C., Fairhead, C., Fok, H., Folegatti, P.M., Fowler, J., Gibbs, C., Goodman, A.L., Jenkin, D., Jones, M., Makinson, R., Marchevsky, N.G., Mujadidi, Y.F., Nguyen, H., Parolini, L., Petersen, C., Plested, E., Pollock, K.M., Ramasamy, M.N., Rhead, S., Robinson, H., Robinson, N., Rongkard, P., Ryan, F., Serrano, S., Tipoe, T., Voysey, M., Waters, A., Zacharopoulou, P., Barnes, E., Dunachie, S., Goulder, P., Klenerman, P., Screaton, G.R., Winston, A., Hill, A.V.S., Gilbert, S.C., Pollard, A.J., Fidler, S., Fox, J., Lambe, T., Watson, M.E.E., Song, R., Cicconi, P., Minassian, A.M., Bibi, S., Kerridge, S., Singh, N., Green, C.M., Douglas, A.D., Lawrie, A.M., Clutterbuck, E.A.: Safety and immunogenicity of the ChAdOx1 nCoV-19 (AZD1222) vaccine against SARS-CoV-2 in HIV infection: a single-arm substudy of a phase 2/3 clinical trial. The Lancet HIV 8(8), 474–485 (2021). 10.1016/S2352-3018(21)00103-X

[20] Brumme, Z.L., Mwimanzi, F., Lapointe, H.R., Cheung, P.K., Sang, Y., Duncan, M.C., Yaseen, F., Agafitei, O., Ennis, S., Ng, K., Basra, S., Lim, L.Y., Kalikawe, R., Speckmaier, S., Moran-Garcia, N., Young, L., Ali, H., Ganase, B., Umviligihozo, G., Harrison Omondi, F., Atkinson, K., Sudderuddin, H., Toy, J., Sereda, P., Burns, L., Costiniuk, C.T., Cooper, C., Anis, A.H., Leung, V., Holmes, D., DeMarco, M.L., Simons, J., Hedgcock, M., Romney, M.G., Barrios, R., Guillemi, S., Brumme, C.J., Pantophlet, R., Montaner, J.S.G., Niikura, M., Harris, M., Hull, M., Brockman, M.A.: Humoral immune responses to COVID-19 vaccination in people living with HIV receiving suppressive antiretroviral therapy. npj Vaccines 7(1) (2022). 10.1038/s41541-022-00452-6

[21] Noe, S., Ochana, N., Wiese, C., Schabaz, F., Von Krosigk, A., Heldwein, S., Rasshofer, R., Wolf, E., Jonsson-Oldenbuettel, C.: Humoral response to SARS-CoV-2 vaccines in people living with HIV. Infection 50(3), 617–623 (2022). 10.1007/s15010-021-01721-7

[22] Knudsen, M.L., Nielsen, S.D., Heftdal, L.D.: Immune responses to mRNA-based vaccines given as a third COVID-19 vaccine dose in people living with HIV—a literature review. Apmis 132(4), 236–244 (2024). 10.1111/apm.13379

[23] Cascarano, A., Mur-Petit, J., Hernández-González, J., Camacho, M., de Toro Eadie, N., Gkontra, P., Chadeau-Hyam, M., Vitrià, J., Lekadir, K.: Machine and Deep Learning for Longitudinal Biomedical Data: a Review of Methods and Applications vol. 56, pp. 1711– 1771. Springer, ??? (2023). 10.1007/s10462-023-10561-w. https://doi.org/10.1007/s10462-023-10561-w

[24] Goronzy, J.J., Weyand, C.M.: Understanding immunosenescence to improve responses to vaccines. Nature Immunology 14(5), 428–436 (2013). 10.1038/ni.2588

[25] Denisko, D., Hoffman, M.M.: Classification and interaction in random forests. Proceedings of the National Academy of Sciences 115(8), 1690– 1692 (2018)

[26] Kumar, S.S., Shaikh, T.: Empirical evaluation of the performance of feature selection approaches on random forest. In: 2017 International Conference on Computer and Applications (ICCA), pp. 227–231. IEEE,(2017)

[27] Van der Maaten, L., Hinton, G.: Visualizing data using t-SNE. Journal of machine learning research 9(11) (2008)

[28] Tuv, E., Borisov, A., Runger, G., Torkkola, K.: Feature selection with ensembles, artificial variables, and redundancy elimination. The Journal of Machine Learning Research 10, 1341–1366 (2009)

[29] Halladin-Dabrowska, A., Kania, A., Kopeć, D.: The t-SNE algorithm as a tool to improve the quality of reference data used in accurate mapping of heterogeneous non-forest vegetation. Remote Sensing 12(1) (2020). 10.3390/RS12010039

[30] Moss, P.: The T cell immune response against SARS-CoV-2. Nature Immunology 23(2), 186–193 (2022). 10.1038/s41590-021-01122-w

[31] Philip, S., S., K.C., Patrick, B., L., C.S.L., Adrian, P., Freda, Q., Melanie, D.-B., R., T.T., Geneviève, M., Monica, D.R., R., A.C., Marc-André, L., Justin, M., Thomas, M., Ryan, L., Erik, M., Salma, S.-M., Yixiao, C.E., Nimitha, P., Anjali, P., Quinn, d.L.K., M., B.J., Alyson, T., Karen, C., Vitaliy, M., Yun, Y.F., Allison, M., Sharon, S., Anne-Claude, G., M., H.J., Mario, O.: mRNA vaccine-induced SARS-CoV-2 spike-specific IFN-*γ* and IL-2 T-cell responses are predictive of serological neutralization and are transiently enhanced by pre-existing cross-reactive immunity. Journal of Virology 0(0), 01685–24 (2025). 10.1128/jvi.01685-24

[32] Lin, J., Law, R., Korosec, C.S., Zhou, C., Koh, H., Ghaemi, S., Samaan, P., Kiang Ooi, H., Matveev, V., Yue, F., Gingras, A.-C., Estacio, A., Buchholz, M., Cheatley, P.L., Mohammadi, A., Kaul, R., Pavinski, K., Mubareka, S., McGeer, A.J., Leis, J.A., Heffernan, J.M., Ostrowski, M.: Longitudinal Assessment of SARS-CoV-2-Specific T Cell Cytokine-Producing Responses for 1 Year Reveals Persistence of Multicytokine Proliferative Responses, with Greater Immunity Associated with Disease Severity. Journal of Virology 96(13), 00509–22 (2022). 10.1128/jvi.00509-22

[33] Baduashvili, A., Guyatt, G., Evans, A.T.: ROC Anatomy—Getting the Most Out of Your Diagnostic Test. Journal of General Internal Medicine 34(9), 1892–1898 (2019). 10.1007/s11606-019-05125-0

[34] Young, C., Brink, R.: The unique biology of germinal center B cells. Immunity 54(8), 1652–1664 (2021). 10.1016/j.immuni.2021.07.015

[35] Good-Jacobson, K.L., Shlomchik, M.J.: Plasticity and Heterogeneity in the Generation of Memory B Cells and Long-Lived Plasma Cells: The Influence of Germinal Center Interactions and Dynamics. The Journal of Immunology 185(6), 3117–3125 (2010). 10.4049/jimmunol.1001155

[36] Janeway, C., Travers, P., Walport, M., Shlomchik, M.: Immunobiology: the Immune System in Health and Disease vol. 2. Garland Pub. New York, ??? (2001)

[37] Mei, H.E., Yoshida, T., Sime, W., Hiepe, F., Thiele, K., Manz, R.A., Radbruch, A., Dörner, T.: Blood-borne human plasma cells in steady state are derived from mucosal immune responses. Blood 113(11), 2461–2469 (2009). 10.1182/blood-2008-04-153544

[38] Wang, Z., Lorenzi, J.C.C., Muecksch, F., Finkin, S., Viant, C., Gaebler, C., Cipolla, M., Hoffmann, H.-H., Oliveira, T.Y., Oren, D.A., Ramos, V., Nogueira, L., Michailidis, E., Robbiani, D.F., Gazumyan, A., Rice, C.M., Hatziioannou, T., Bieniasz, P.D., Caskey, M., Nussenzweig, M.C.: Enhanced SARS-CoV-2 neutralization by dimeric IgA. Science Translational Medicine 13(577), 1555 (2021). 10.1126/scitranslmed.abf1555

[39] Sheikh-Mohamed, S., Isho, B., Chao, G.Y.C., Zuo, M., Cohen, C., Lustig, Y., Nahass, G.R., Salomon-Shulman, R.E., Blacker, G., Fazel-Zarandi, M., Rathod, B., Colwill, K., Jamal, A., Li, Z., de Launay, K.Q., Takaoka, A., Garnham-Takaoka, J., Patel, A., Fahim, C., Paterson, A., Li, A.X., Haq, N., Barati, S., Gilbert, L., Green, K., Mozafarihashjin, M., Samaan, P., Budylowski, P., Siqueira, W.L., Mubareka, S., Ostrowski, M., Rini, J.M., Rojas, O.L., Weissman, I.L., Tal, M.C., McGeer, A., Regev-Yochay, G., Straus, S., Gingras, A.-C., Gommerman, J.L.: Systemic and mucosal IgA responses are variably induced in response to SARS-CoV-2 mRNA vaccination and are associated with protection against subsequent infection. Mucosal Immunology 15(5), 799–808 (2022). 10.1038/s41385-022-00511-0

[40] Hel, Z., McGhee, J.R., Mestecky, J.: HIV infection: first battle decides the war. Trends in Immunology 27(6), 274–281 (2006). 10.1016/j.it.2006.04.007

[41] Xu, W., Santini, P.A., Sullivan, J.S., He, B., Shan, M., Ball, S.C., Dyer, W.B., Ketas, T.J., Chadburn, A., Cohen-Gould, L., Knowles, D.M., Chiu, A., Sanders, R.W., Chen, K., Cerutti, A.: HIV-1 evades virus-specific IgG2 and IgA responses by targeting systemic and intestinal B cells via long-range intercellular conduits. Nature Immunology 10(9), 1008–1017 (2009). 10.1038/ni.1753

[42] Allers, K., Puyskens, A., Epple, H.J., Schürmann, D., Hofmann, J., Moos, V., Schneider, T.: The effect of timing of antiretroviral therapy on CD4+ T-cell reconstitution in the intestine of HIV-infected patients. Mucosal Immunology 9(1), 265–274 (2016). 10.1038/mi.2015.58

[43] Tincati, C., Douek, D.C., Marchetti, G.: Gut barrier structure, mucosal immunity and intestinal microbiota in the pathogenesis and treatment of HIV infection. AIDS Research and Therapy 13(1), 1–11 (2016). 10.1186/s12981-016-0103-1

[44] Thwaites, R.S., Uruchurtu, A.S.S., Negri, V.A., Cole, M.E., Singh, N., Poshai, N., Jackson, D., Hoschler, K., Baker, T., Scott, I.C., Ros, X.R., Cohen, E.S., Zambon, M., Pollock, K.M., Hansel, T.T., Openshaw, P.J.M.: Early mucosal events promote distinct mucosal and systemic antibody responses to live attenuated influenza vaccine. Nature Communications 14(1), 1–14 (2023). 10.1038/s41467-023-43842-7

[45] Buckner, C.M., Moir, S., Ho, J., Wang, W., Posada, J.G., Kardava, L., Funk, E.K., Nelson, A.K., Li, Y., Chun, T.-W., Fauci, A.S.: Characterization of Plasmablasts in the Blood of HIV-Infected Viremic Individuals: Evidence for Nonspecific Immune Activation. Journal of Virology 87(10), 5800–5811 (2013). 10.1128/jvi.00094-13

[46] Qiao, X., He, B., Chiu, A., Knowles, D.M., Chadburn, A., Cerutti, A.: Human immunodeficiency virus 1 Nef suppresses CD40-dependent immunoglobulin class switching in bystander B cells. Nature Immunology 7(3), 302–310 (2006). 10.1038/ni1302

[47] Chen, R.J., Lu, M.Y., Chen, T.Y., Williamson, D.F.K., Mahmood, F.: Synthetic data in machine learning for medicine and healthcare. Nature Biomedical Engineering 5(6), 493–497 (2021). 10.1038/s41551-021-00751-8

[48] Giuffrè, M., Shung, D.L.: Harnessing the power of synthetic data in healthcare: innovation, application, and privacy. npj Digital Medicine 6(1), 1–8 (2023). 10.1038/s41746-023-00927-3

[49] Iwasaki, A., Medzhitov, R.: Control of adaptive immunity by the innate immune system. Nature immunology 16(4), 343–353 (2015). 10.1038/ni.3123

[50] Korosec, C.S., Betti, M.I., David, W., Ooi, H.K., Moyles, I.R., Wahl, L.M., Heffernan, J.M.: Multiple cohort study of hospitalized SARS-CoV-2 in-host infection dynamics: Parameter estimates, identifiability, sensitivity and the eclipse phase profile. Journal of theoretical biology 564, 111449 (2023) [arXiv:2022.06.20.22276662](https://arXiv.org/abs/2022.06.20.22276662). 10.1016/j.jtbi.2023.111449

[51] Farhang-sardroodi, S., Korosec, C.S., Gholami, S., Craig, M., Moyles, I.R., Ghaemi, M.S., Ooi, H.K., Heffernan, J.M.: Analysis of Host Immunological Response of Adenovirus-Based COVID-19 Vaccines. Vaccines 9(8), 861 (2021)

[52] Moyles, I.R., Korosec, C.S., Heffernan, J.M.: Determination of significant immunological timescales from mRNA-LNP-based vaccines in humans. Journal of Mathematical Biology 86(86), 1–41 (2023) [arXiv:2022.07.25.22278031](https://arXiv.org/abs/2022.07.25.22278031). 10.1007/s00285-023-01919-3

[53] Korosec, C.S., Farhang-Sardroodi, S., Dick, D.W., Gholami, S., Ghaemi, M.S., Moyles, I.R., Craig, M., Ooi, H.K., Heffernan, J.M.: Long-term durability of immune responses to the BNT162b2 and mRNA-1273 vaccines based on dosage, age and sex. Scientific Reports 12(1), 21232 (2022) arXiv:/doi.org/10.1101/2021.10.13.21264957 [https:]. 10.1038/s41598-022-25134-0

[54] Korosec, C.S., Dick, D.W., Moyles, I.R., Watmough, J.: SARS-CoV-2 booster vaccine dose significantly extends humoral immune response halflife beyond the primary series. Scientific Reports 14(1), 8426 (2024). 10.1038/s41598-024-58811-3

[55] Gholami, S., Korosec, C.S., Farhang-Sardroodi, S., Dick, D.W., Craig, M., Ghaemi, M.S., Ooi, H.K., Heffernan, J.M.: A mathematical model of protein subunits COVID-19 vaccines. Mathematical Biosciences 358, 108970 (2023). 10.1016/J.MBS.2023.108970

[56] Banuet-Martinez, M., Yang, Y., Jafari, B., Kaur, A., Butt, Z.A., Chen, H.H., Yanushkevich, S., Moyles, I.R., Heffernan, J.M., Korosec, C.S.: Monkeypox: A review of epidemiological modelling studies and how modelling has led to mechanistic insight. Epidemiology and Infection 151, 121 (2023). 10.1017/S0950268823000791

[57] Stelzer, I.A., Ghaemi, M.S., Han, X., Ando, K., Hédou, J.J., Feyaerts, D., Peterson, L.S., Rumer, K.K., Tsai, E.S., Ganio, E.A., Gaudillière, D.K., Tsai, A.S., Choisy, B., Gaigne, L.P., Verdonk, F., Jacobsen, D., Gavasso, S., Traber, G.M., Ellenberger, M., Stanley, N., Becker, M., Culos, A., Fallahzadeh, R., Wong, R.J., Darmstadt, G.L., Druzin, M.L., Winn, V.D., Gibbs, R.S., Ling, X.B., Sylvester, K., Carvalho, B., Snyder, M.P., Shaw, G.M., Stevenson, D.K., Contrepois, K., Angst, M.S., Aghaeepour, N., Gaudillière, B.: Integrated trajectories of the maternal metabolome, proteome, and immunome predict labor onset. Science Translational Medicine 13(592) (2021). 10.1126/scitranslmed.abd9898

[58] Ghaemi, M.S., DiGiulio, D.B., Contrepois, K., Callahan, B., Ngo, T.T.M., Lee-Mcmullen, B., Lehallier, B., Robaczewska, A., McIlwain, D., Rosenberg-Hasson, Y., Wong, R.J., Quaintance, C., Culos, A., Stanley, N., Tanada, A., Tsai, A., Gaudilliere, D., Ganio, E., Han, X., Ando, K., McNeil, L., Tingle, M., Wise, P., Maric, I., Sirota, M., Wyss-Coray, T., Winn, V.D., Druzin, M.L., Gibbs, R., Darmstadt, G.L., Lewis, D.B., Partovi Nia, V., Agard, B., Tibshirani, R., Nolan, G., Snyder, M.P., Relman, D.A., Quake, S.R., Shaw, G.M., Stevenson, D.K., Angst, M.S., Gaudilliere, B., Aghaeepour, N.: Multiomics modeling of the immunome, transcriptome, microbiome, proteome and metabolome adaptations during human pregnancy. Bioinformatics 35(1), 95–103 (2019). 10.1093/bioinformatics/bty537

[59] Aghaeepour, N., Ganio, E.A., Mcilwain, D., Tsai, A.S., Tingle, M., Van Gassen, S., Gaudilliere, D.K., Baca, Q., McNeil, L., Okada, R., Ghaemi, M.S., Furman, D., Wong, R.J., Winn, V.D., Druzin, M.L., El-Sayed, Y.Y., Quaintance, C., Gibbs, R., Darmstadt, G.L., Shaw, G.M., Stevenson, D.K., Tibshirani, R., Nolan, G.P., Lewis, D.B., Angst, M.S., Gaudilliere, B.: An immune clock of human pregnancy. Science Immunology 2(15), 1–11 (2017). 10.1126/sciimmunol.aan2946

[60] Culos, A., Tsai, A.S., Stanley, N., Becker, M., Ghaemi, M.S., McIlwain, D.R., Fallahzadeh, R., Tanada, A., Nassar, H., Espinosa, C., Xenochristou, M., Ganio, E., Peterson, L., Han, X., Stelzer, I.A., Ando, K., Gaudilliere, D., Phongpreecha, T., Marić, I., Chang, A.L., Shaw, G.M., Stevenson, D.K., Bendall, S., Davis, K.L., Fantl, W., Nolan, G.P., Hastie, T., Tibshirani, R., Angst, M.S., Gaudilliere, B., Aghaeepour, N.: Integration of mechanistic immunological knowledge into a machine learning pipeline improves predictions. Nature Machine Intelligence 2(10), 619–628 (2020). 10.1038/s42256-020-00232-8

[61] Pardi, N., Krammer, F.: mRNA vaccines for infectious diseases — advances, challenges and opportunities. Nature Reviews Drug Discovery (2024). 10.1038/s41573-024-01042-y

[62] Colwill, K., Galipeau, Y., Stuible, M., Gervais, C., Arnold, C., Rathod, B., Abe, K.T., Wang, J.H., Pasculescu, A., Maltseva, M., Rocheleau, L., Pelchat, M., Fazel-Zarandi, M., Iskilova, M., Barrios-Rodiles, M., Bennett, L., Yau, K., Cholette, F., Mesa, C., Li, A.X., Paterson, A., Hladunewich, M.A., Goodwin, P.J., Wrana, J.L., Drews, S.J., Mubareka, S., McGeer, A.J., Kim, J., Langlois, M.-A., Gingras, A.-C., Durocher, Y.: A scalable serology solution for profiling humoral immune responses to SARS-CoV-2 infection and vaccination. Clinical & Translational Immunology 11(3), 1380 (2022). 10.1002/cti2.1380

[63] Isho, B., Abe, K.T., Zuo, M., Jamal, A.J., Rathod, B., Wang, J.H., Li, Z., Chao, G., Rojas, O.L., Bang, Y.M., Pu, A., Christie-Holmes, N., Gervais, C., Ceccarelli, D., Samavarchi-Tehrani, P., Guvenc, F., Budylowski, P., Li, A., Paterson, A., Yue, F.Y., Marin, L.M., Caldwell, L., Wrana, J.L., Colwill, K., Sicheri, F., Mubareka, S., Gray-Owen, S.D., Drews, S.J., Siqueira, W.L., Barrios-Rodiles, M., Ostrowski, M., Rini, J.M., Durocher, Y., McGeer, A.J., Gommerman, J.L., Gingras, A.-C.: Persistence of serum and saliva antibody responses to SARS-CoV-2 spike antigens in COVID-19 patients. Science Immunology 5(52), 5511 (2020). 10.1126/sciimmunol.abe5511

[64] Liang, W., Liang, H., Ou, L., Chen, B., Chen, A., Li, C., Li, Y., Guan, W., Sang, L., Lu, J., Xu, Y., Chen, G., Guo, H., Guo, J., Chen, Z., Zhao, Y., Li, S., Zhang, N., Zhong, N., He, J.: Development and validation of a clinical risk score to predict the occurrence of critical illness in hospitalized patients with COVID-19. JAMA Internal Medicine 180(8), 1081–1089 (2020). 10.1001/jamainternmed.2020.2033

[65] Austin, P.C., White, I.R., Lee, D.S., van Buuren, S.: Missing Data in Clinical Research: A Tutorial on Multiple Imputation. Canadian Journal of Cardiology 37(9), 1322–1331 (2021). 10.1016/j.cjca.2020.11.010

[66] Stekhoven, D.J., Bühlmann, P.: Missforest-Non-parametric missing value imputation for mixed-type data. Bioinformatics 28(1), 112–118 (2012) [arXiv:1105.0828](https://arXiv.org/abs/1105.0828). 10.1093/bioinformatics/btr597

[67] Waljee, A.K., Mukherjee, A., Singal, A.G., Zhang, Y., Warren, J., Balis, U., Marrero, J., Zhu, J., Higgins, P.D.R.: Comparison of imputation methods for missing laboratory data in medicine. BMJ Open 3(8), 1–7 (2013). 10.1136/bmjopen-2013-002847

[68] Khademi, A.: Flexible imputation of missing data 2nd edition. Journal of Statistical Software 93(April), 1–4 (2020). 10.18637/jss.v093.b01

[69] Tiwaskar, S., Rashid, M., Gokhale, P.: Impact of machine learningbased imputation techniques on medical datasets- a comparative analysis. Multimedia Tools and Applications (2024). 10.1007/s11042-024-19103-0

[70] Krijthe, J.: R Package ‘Rtsne’. CRAN (2023). https://cran.r-project.org/web/packages/Rtsne/Rtsne.pdf

[71] Breiman, L.: Random Forests. Machine Learning 45(1), 5–32 (2001). 10.1023/A:1010933404324

