## Supplementary material for "Machine Learning Reveals Distinct Immunogenic Signatures of Th1 Imprinting in ART-Treated Individuals with HIV Following Repeated SARS-CoV-2 Vaccination"

March 18, 2025

AUTHORS

Corresponding author emails:

### Contents

**S1 RF classification probabilities** **2**

S1.1 Synthetic Data Analysis . . . . . 7

**S2 Imputation analysis** **11**

### S1 RF classification probabilities

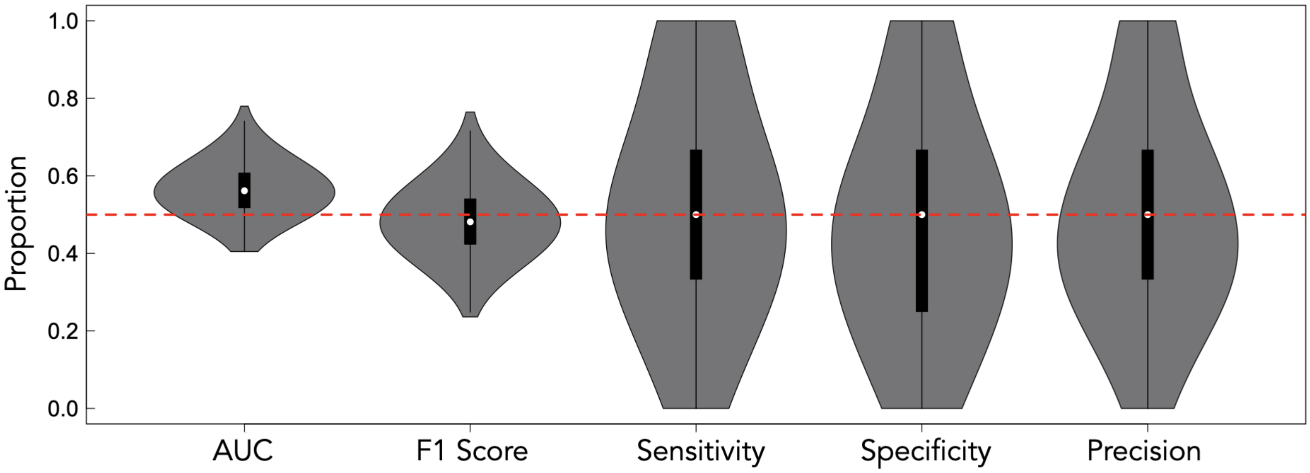


Figure S1: Model performance with randomized labels using all 63 features. RF was carried out similarly to as described in the methods section with K-fold CV, however, the labels were randomized. Provided are the AUC, F1 Score, Sensitivity, Specificity, and Precision, respectively, which can be compared to the true model performance metrics provided in Fig 1E. A red dashed line at 0.5 is shown for comparison

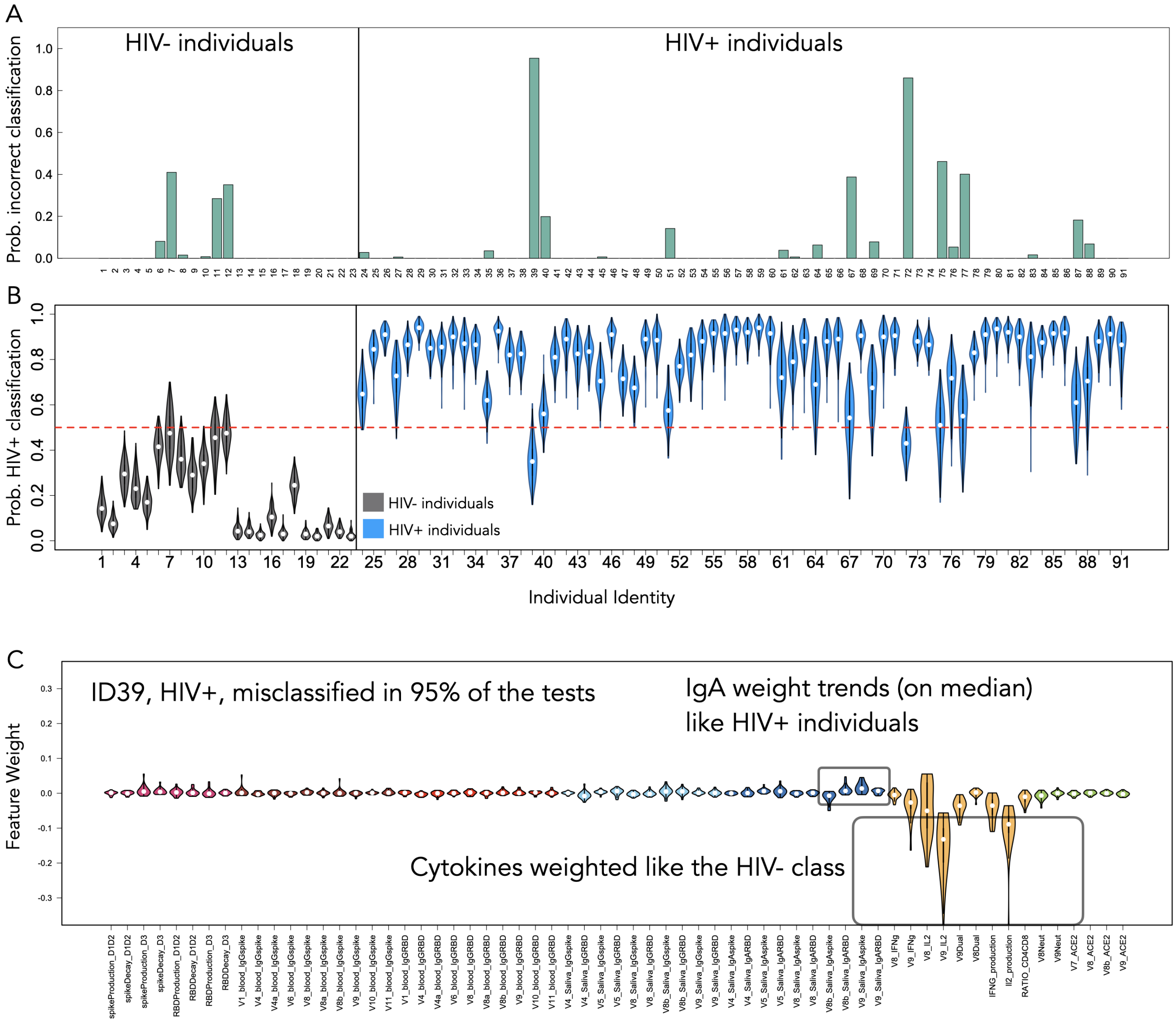


Figure S2: A) Probability that the RF model will incorrectly classify the individual, for all individuals in the full-feature RF models. B) Violin plots for the distributions of HIV+ classifications for all individuals. C) Example feature weight distributions for ID39, who tends to be misclassified as HIV- when they are known to be HIV+.

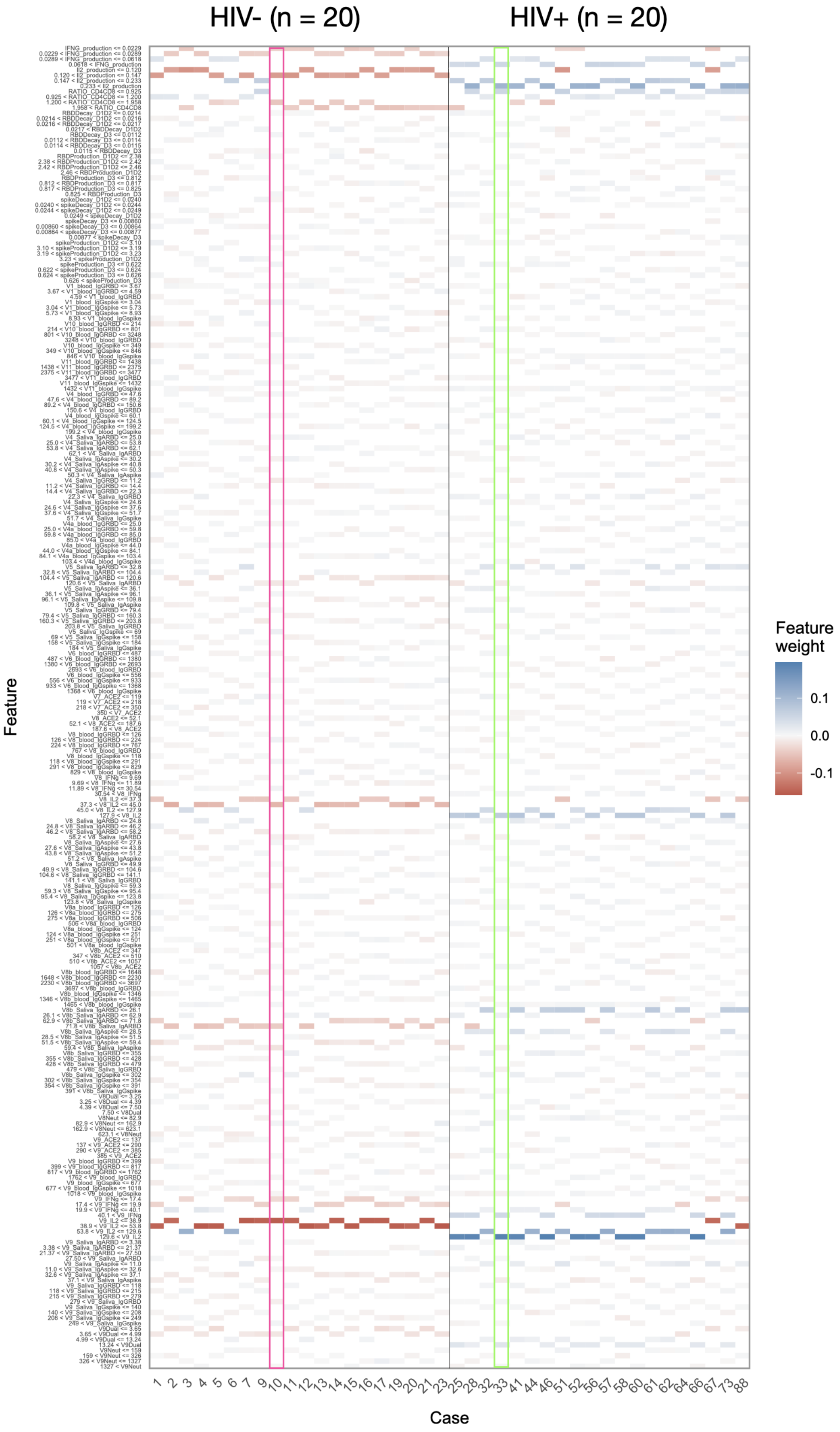


Figure S3: An example of the RF training landscape for a single RF model, IDs 10 and 33 are randomly highlighted, with their corresponding individual feature distributions shown in Fig. S4. This RF model produces an AUC of 1.0.

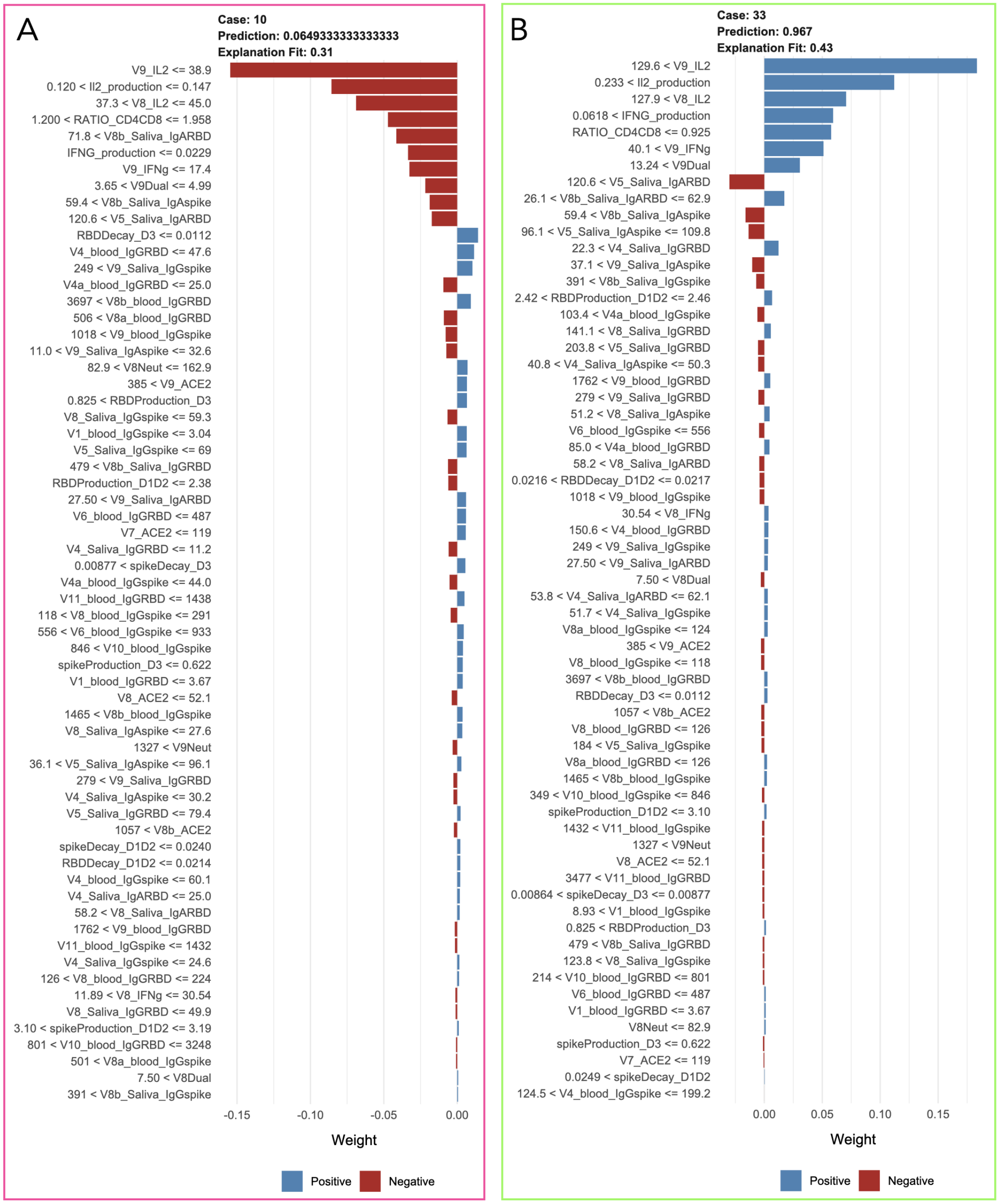


Figure S4: Individual feature weights for IDs 10 and ID 33 for a single RF model trained on all 64 features from the model landscape shown in Fig. S3.

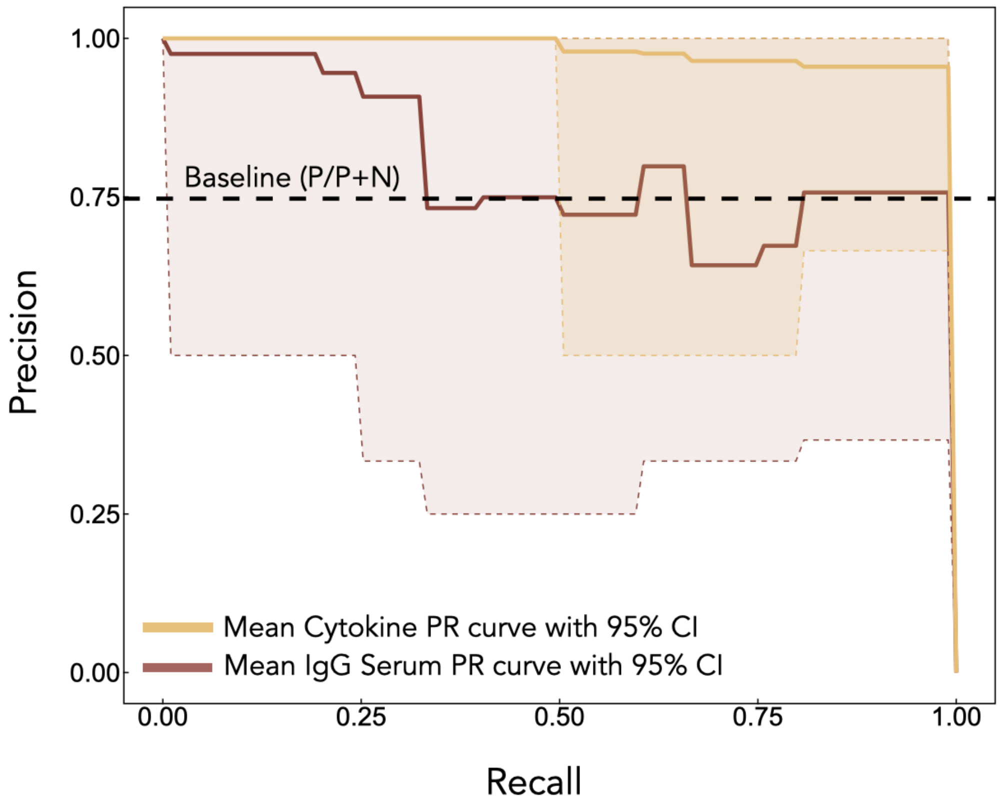


Figure S5: Mean PR curves for RF models trained on just the cytokine features (yellow) and serum data (red). Shaded regions represent 95% confidence intervals. The baseline (dashed line) is calculated by computing the ratio $P/(P+N)$ which is the ratio of positives and negatives in the full data set (not the downsampled balanced ratio of 1:1 used for training, but the representative ratio used for testing). In accordance with intuition gained from the mean ROC curves (Fig. 3E), the cytokine features result in a near-perfect classifier while serum features result in approximately baseline (random) performance.

#### S1.1 Synthetic Data Analysis

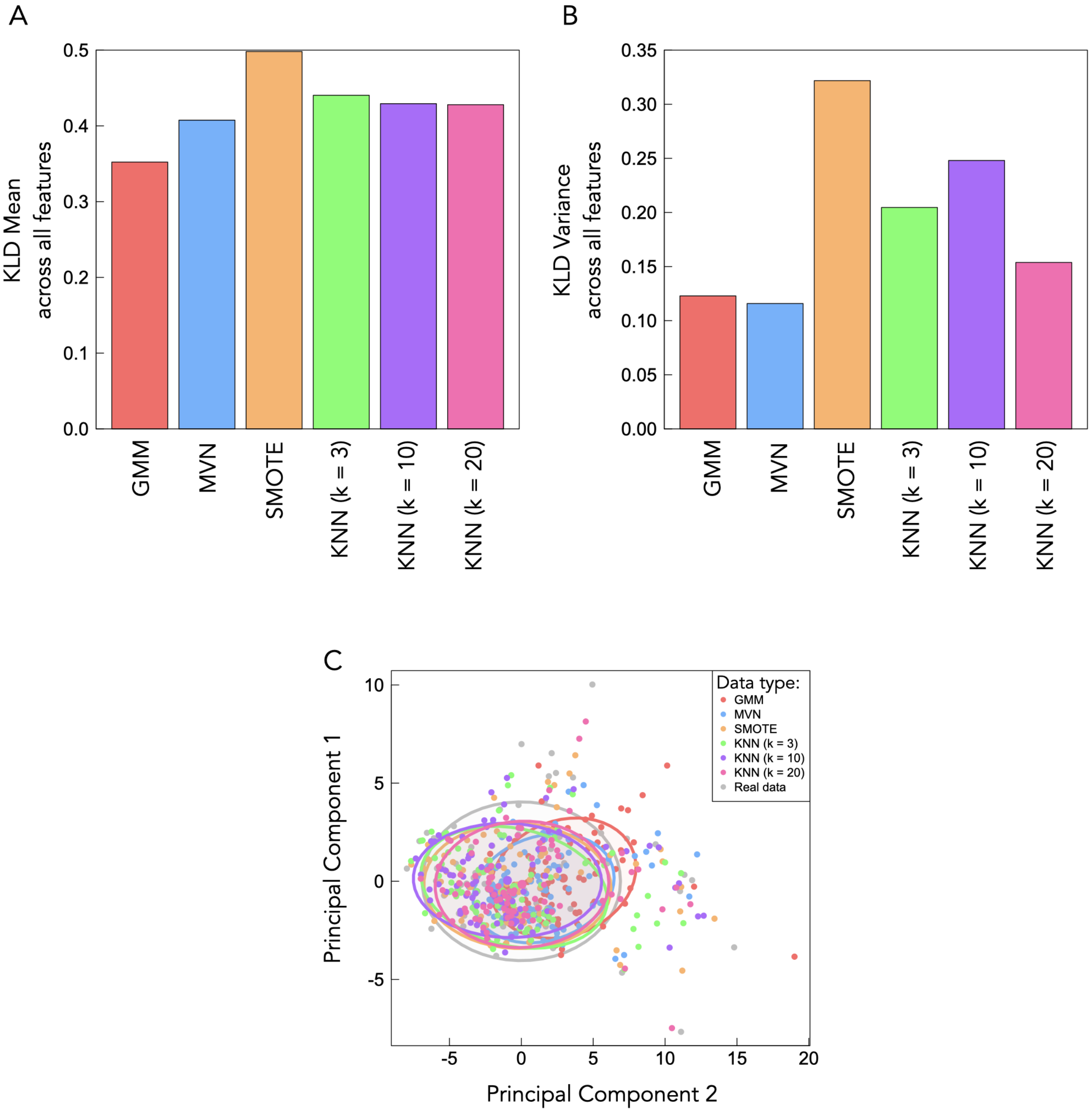


Figure S6: A) Mean KLD is computed across all features between each respective synthetic data set and the real data. B) The variance in KLD is computed across all features between each respective synthetic data set and the real data. C) PCA analysis is displayed whereby the principal components from the real data are projected onto each respective principal components of the synthetic data. Ellipses are first standard deviation. Here, all synthetic data approaches are reasonably approximate the actual data and no meaningful cluster separation is found.

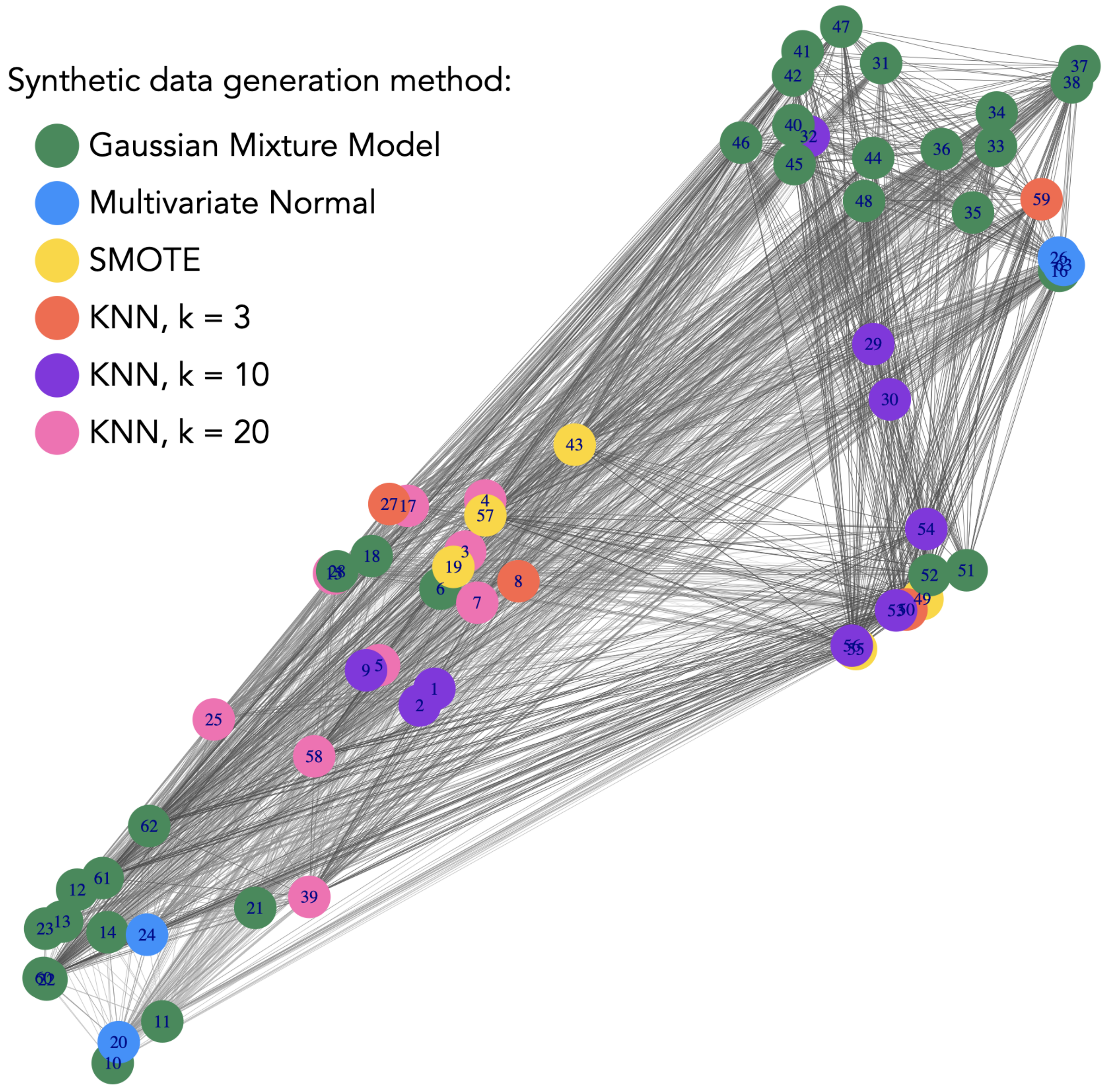


Figure S7: t-SNE plot shown with the same layout as in the maintext, however, colours here correspond to the synthetic data feature method that minimized the KLD for that specific feature.

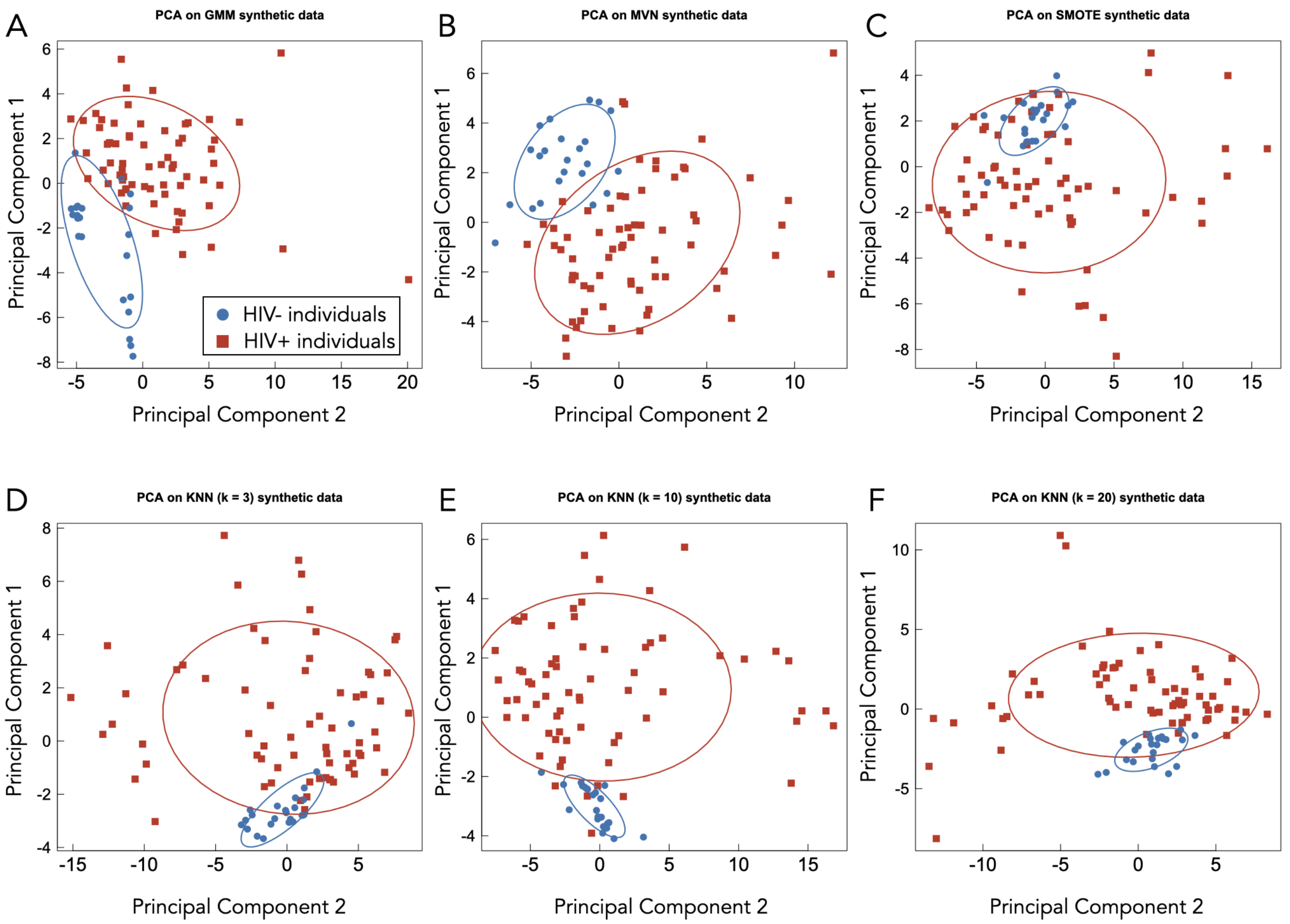


Figure S8: PCA on synthetically generated data. Panels A-F display the results of PCA performed on the synthetically generated data. Panels are ordered by GMM, MVN, SMOTE, KNN (k = 3), KNN (k = 10), and KNN (k = 20), for panels A, B, C, D, E, and F, respectively. SMOTE, despite having the highest mean KLD, is the only technique whereby the control class subclusters within the first standard deviation ellipse of the target class, which is similar structural behaviour observed by the actual data (Fig.1C). SMOTE is the only supervised method used to generate synthetic data in this work.

#### S2 Imputation analysis

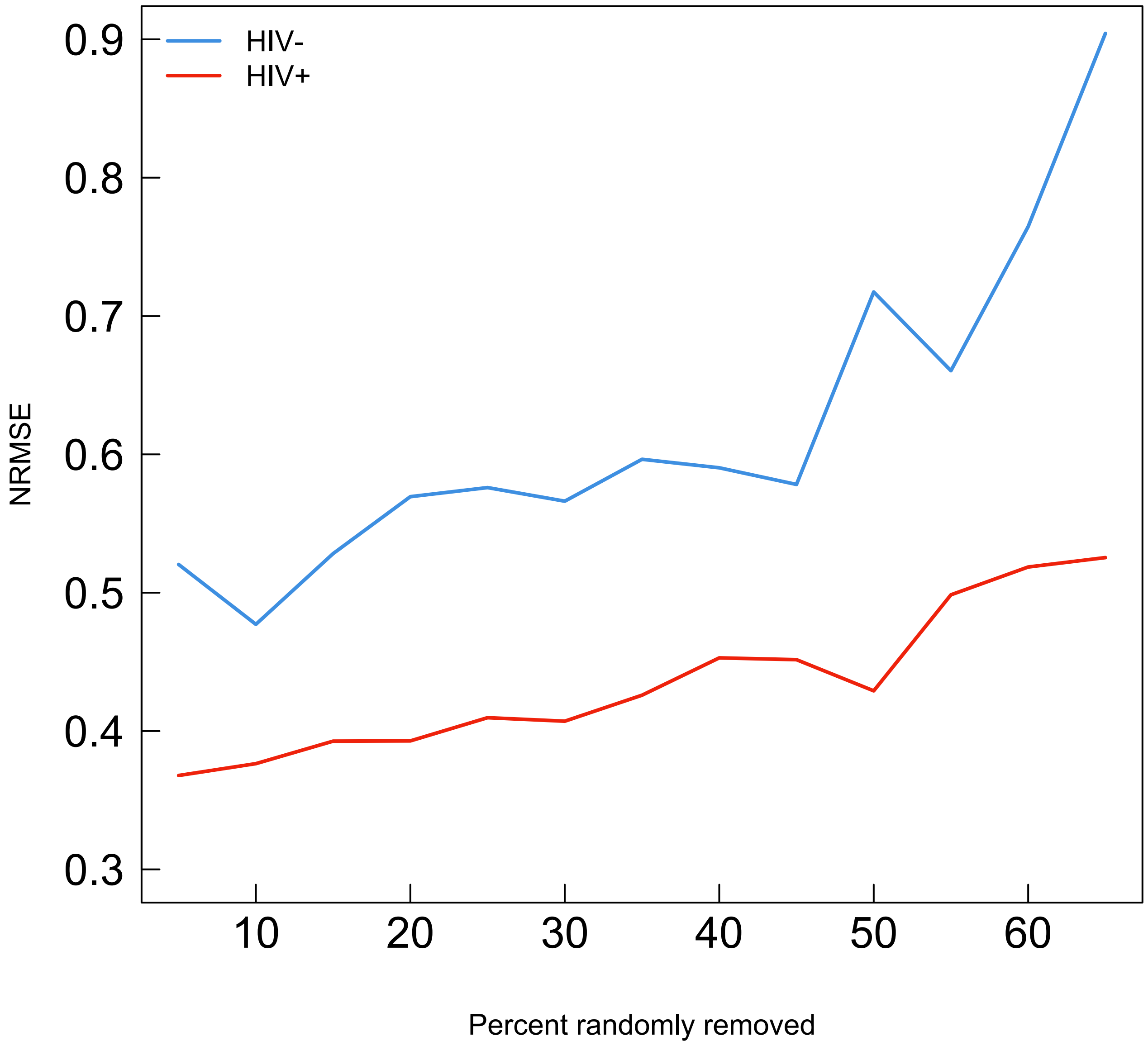


Figure S9: Normalized Root Mean Squared Error (NRMSE) of imputed features as a function of increasing missingness. NRMSE was computed separately for HIV-negative (blue) and HIV-positive (red) groups, with each point representing the average NRMSE across all features at a given level of missing data. As expected, NRMSE increases gradually as the proportion of missing data increases, reflecting the increasing uncertainty in imputed values. However, the shallow slope suggests that the imputation method remains robust across varying levels of missingness, preserving data integrity even at higher rates of removal.

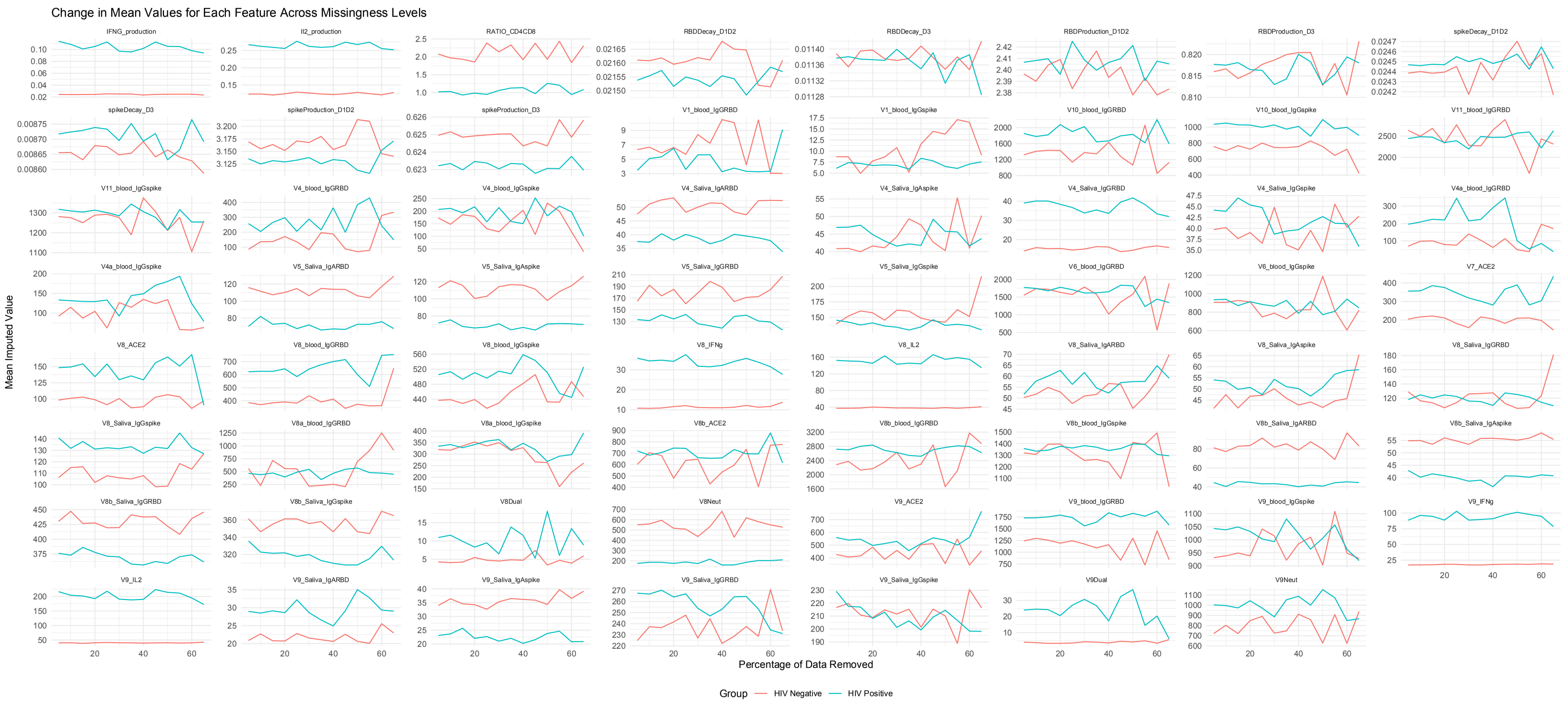


Figure S10: Mean values of the imputed features remains stable across increasing levels of missingness. Each panel represents a single feature, with the x-axis indicating the percentage of data removed prior to imputation, and the y-axis representing the mean of the imputed values. Imputation was performed separately for HIV-negative (blue) and HIV-positive (red) groups as described in the methods section. The relative stability of these metrics suggests that the imputation approach preserves the underlying data structure, even as missingness increases.

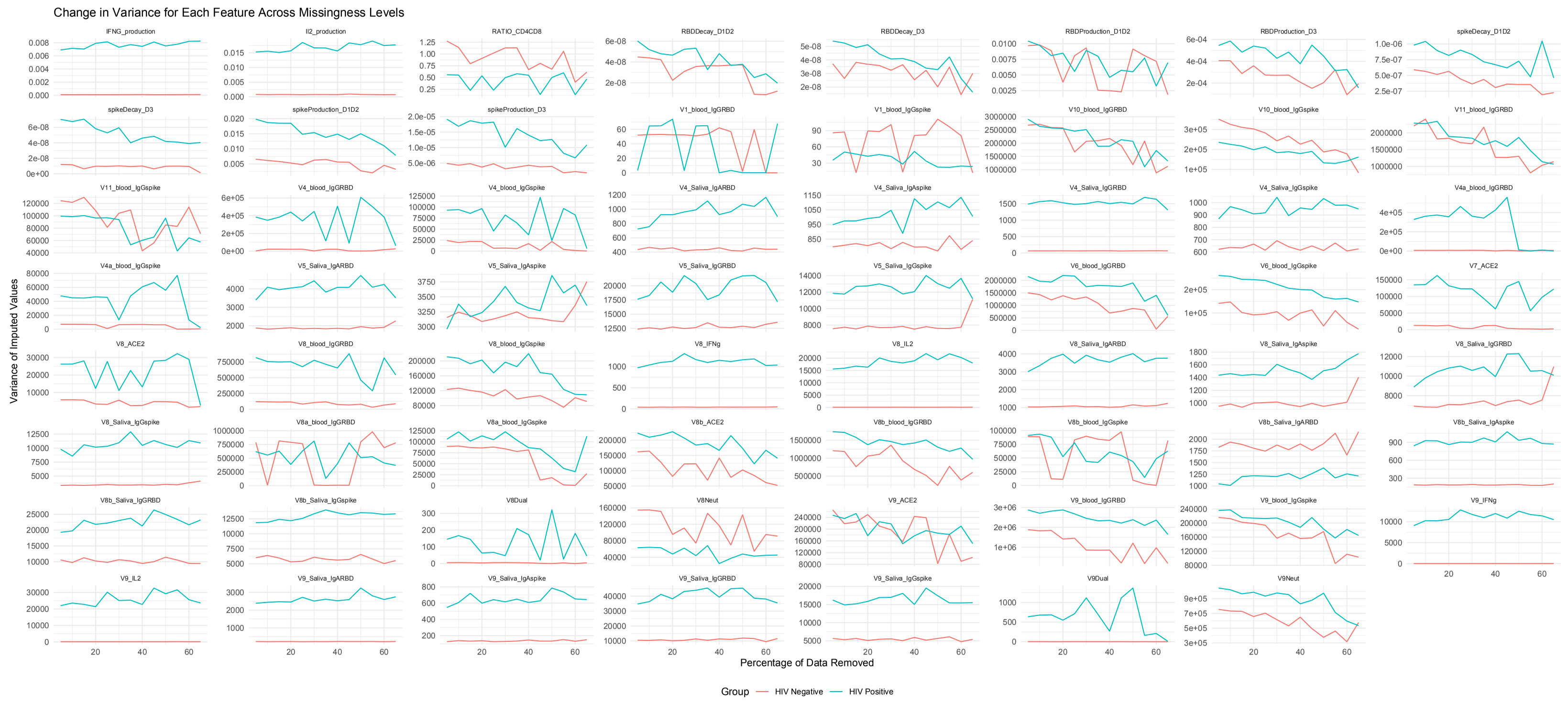


Figure S11: Variance of the imputed features remains stable across increasing levels of missingness. Each panel represents a single feature, with the x-axis indicating the percentage of data removed prior to imputation, and the y-axis representing the mean of the imputed values. Imputation was performed separately for HIV-negative (blue) and HIV-positive (red) groups as described in the methods section. The relative stability of these metrics suggests that the imputation approach preserves the underlying data structure, even as missingness increases.
